# Astrocytes drive divergent metabolic gene expression in humans and chimpanzees

**DOI:** 10.1101/2020.11.09.374835

**Authors:** Trisha M. Zintel, Jason Pizzollo, Christopher G. Claypool, Courtney C. Babbitt

**Affiliations:** Department of Biology, University of Massachusetts Amherst, Amherst, MA; Molecular and Cellular Biology Graduate Program, University of Massachusetts Amherst, Amherst, MA; Organismic and Evolutionary Biology, University of Massachusetts Amherst, Amherst, MA

**Author notes:** Corresponding author: Courtney C. Babbitt, Ph.D., Dept. of Biology, 611 North Pleasant St., Amherst, MA, 01003, USA.

**Keywords:** astrocytes, neurons, evolution, metabolism, genomics, brain

## Abstract

The human brain utilizes ∼ 20% of all of the body’s metabolic resources, while chimpanzee brains use less than 10%. Although previous work shows significant differences in metabolic gene expression between the brains of primates, we have yet to fully resolve the contribution of distinct brain cell types. To investigate cell-type specific interspecies differences in brain gene expression, we conducted RNA-Seq on neural progenitor cells (NPCs), neurons, and astrocytes generated from induced pluripotent stem cells (iPSCs) from humans and chimpanzees. Interspecies differential expression (DE) analyses revealed that twice as many genes exhibit DE in astrocytes (12.2% of all genes expressed) than neurons (5.8%). Pathway enrichment analyses determined that astrocytes, rather than neurons, diverged in expression of glucose and lactate transmembrane transport, as well as pyruvate processing and oxidative phosphorylation. These findings suggest that astrocytes may have contributed significantly to the evolution of greater brain glucose metabolism with proximity to humans.

## INTRODUCTION

Though primates exhibit widespread variation in many phenotypes, including anatomy, behavior, and cognition, the extent of these phenotypic differences is not substantially larger than differences in genome sequence (A. Varki & Altheide, 2005; N. M. Varki & Varki, 2015). One of those traits that defines primates is a significantly larger brain relative to body size, for which humans exhibit the greatest amount of difference. Within primates, selective differences in the genome can be linked to diet and metabolism, suggesting selection has optimized different metabolic processes in lineage-dependent ways (Babbitt et al., 2010; Babbitt, Warner, Fedrigo, Wall, & Wray, 2011; Bauernfeind et al., 2015; Fagundes et al., 2007; Haygood, Babbitt, Fedrigo, & Wray, 2010; Schaffner et al., 2005; Stringer & Andrews, 1988). The human brain is more energetically costly than that of other primates, utilizing ∼ 20% of all of the body’s metabolic resources, in comparison to non-human primate brains that use less than 10% (Hofman, 1983; Mink, Blumenschine, & Adams, 1981). Importantly, allometry alone does not explain the increase in human brain appropriation of glucose metabolism at this proportion (Karbowski, 2007; Martin, 1981; Yu, Karbowski, Sachdev, & Feng, 2014). There is evidence that sheer increase in neuron number can explain at least part of the energetic demand of the human brain (Herculano-Houzel, 2011). However, interspecies differences in the contribution of metabolism of astrocytes versus neurons to metabolic capacity at the organ level remain largely un-explored.

Many of these changes in brain metabolism have been hypothesized to coincide with other trait changes, particularly those related to shifts in diet known to be important in hominin evolution, such as an increase in meat products, increased quality of food, and agriculture (Aiello & Wheeler, 1995; Babbitt et al., 2011; F. Brown, Harris, Leakey, & Walker, 1985; McHenry, 1992, 1994; Peters, 2007; Shea, 2007). Furthermore, the expensive-tissue hypothesis posits that a trade-off in energy allocation occurred between energetically expensive or storing tissues for the development of a larger, metabolically demanding brain in primates, including a reduction in energetically expensive gut tissue (Aiello & Wheeler, 1995; Pontzer et al., 2014; Stearns, 1992; West, Brown, & Enquist, 2001) or a shift to investment in energy-storing tissue adipose tissue rather than energy-utilizing muscle tissue (Leonard & Robertson, 1994, 1997; Leonard, Robertson, Snodgrass, & Kuzawa, 2003). There is evidence that the higher metabolic costs of the human brain influences the protracted development of body growth rate (Kuzawa et al., 2014). These *in vivo* (whole organism) studies further suggest an important link between evolutionary differences in metabolism and the uniqueness of the primate brain.

Similar to organism-level investigations, there is also molecular evidence supporting the evolution of metabolic processes (e.g. oxidative phosphorylation) in the primate brain with phylogenetic proximity to humans. Metabolism in the brain is critical for neurological function, as it provides cellular energy and critical biomolecules necessary for the complex cellular network characteristic of the brain (Bauernfeind & Babbitt, 2014; A. M. Brown, Wender, & Ransom, 2001; Nelson, Lehninger, & Cox, 2008; Raichle, 2010; Tekkök, Brown, Westenbroek, Pellerin, & Ransom, 2005; Vander Heiden, Cantley, & Thompson, 2009; Vander Heiden et al., 2010). Cellular metabolism involves the breakdown of fuel molecules to produce energy or other molecules through multiple interconnected pathways, including glycolysis, oxidative phosphorylation, and the pentose phosphate pathway. Enrichments for metabolic processes in genes and gene regulatory regions undergoing positive selection is a common thread in gene expression analyses from whole primate brain tissue (Babbitt et al., 2010; Bauernfeind et al., 2015; Haygood et al., 2010; Haygood, Fedrigo, Hanson, Yokoyama, & Wray, 2007; Kosiol et al., 2008; Uddin et al., 2008). Interestingly, there are lineage-dependent differences in the specific pathways enriched in each species (e.g. glucose and carbohydrate metabolism in humans and glycogen and acyl-CoA metabolism in chimpanzees) (Haygood et al., 2007; Kosiol et al., 2008; Uddin et al., 2008). Within anthropoids, genes encoding the subunits of cytochrome c oxidase, the final component of the electron transport chain, show an accelerated rate of evolution in their sequences compared with any other placental mammals (Grossman, Schmidt, Wildman, & Goodman, 2001; Uddin et al., 2008; Wildman, Wu, Goodman, & Grossman, 2002; Wu, Goodman, Lomax, & Grossman, 1997). These molecular changes suggest increased control over the mechanisms that process glucose (Goldberg et al., 2003; Grossman et al., 2001; Grossman, Wildman, Schmidt, & Goodman, 2004; Hüttemann et al., 2012; Uddin et al., 2008). Further understanding the relationship between genetic changes (both in coding and non-coding regulatory portions of the genome) and observed metabolic differences in primates will contribute to a greater understanding of proximate influences on larger evolutionary trends in primates. These findings also highlight a need to investigate not only glucose metabolism and energy production but also that of other macromolecules (e.g. lipids, amino acids, nucleic acids) for a more comprehensive understanding of differences in cellular metabolism in neural cells of primates.

Many previous comparative primate studies using functional genomics have determined significant differences in expression between humans, chimpanzees, and other primate species (primarily, rhesus macaque) (Babbitt et al., 2010; Bakken et al., 2015; Bauernfeind et al., 2015; Blekhman, Oshlack, Chabot, Smyth, & Gilad, 2008; Khaitovich et al., 2004; Konopka et al., 2012; Oldham, Horvath, & Geschwind, 2006). However, as many investigations of primate evolution and humans in particular often are, these studies have been largely limited to utilizing post-humous tissue samples, oftentimes opportunistically obtained. Recent advances in induced pluripotent stem cell (iPSC) technology have allowed for the generation of iPSC-derived mature brain cells and organoids as *in vitro* models of primate brain development. There have been a number of studies using iPSCs to generate brain organoids as an *in vitro* model for brain development (Amiri et al., 2018; Camp et al., 2015; Luo et al., 2016; Velasco et al., 2019), including one comparative study between humans, chimpanzees, and rhesus macaques (Kanton et al., 2019). The use of iPSC-derived samples has shown great promise in understanding brain development and function in greater detail. While monolayer culturing of iPSC-derived cells does not recapitulate the complexity of the primate brain as well as brain organoids, they are far more feasible in both cost and time, and have been used extensively to investigate brain cell-type specific mechanisms of disease (Cho, Yang, Forest, Qian, & Chan, 2019; di Domenico et al., 2019; Penney, Ralvenius, & Tsai, 2019; Zhao et al., 2017).

The findings of interspecies divergence in brain metabolism are intriguing, however, a cell-type specific comparison would more fully inform our understanding of distinct cellular contributions to interspecific differences in neurological function (I. G. Romero et al., 2015). Two of the major cell types in the brain are neurons and astrocytes. Neurons function in neurological processes like cognition and perception largely by transmitting chemical and electrical signals throughout complex cellular networks. However, metabolic programs have been shown to shift as neural progenitor cells (NPCs) differentiate into more mature cell types (Zheng et al., 2016). Non-dividing, mature neurons are known to have very little capacity for specific metabolic processes (e.g. glycolysis) and rely on metabolite shuttling from another cell type, astrocytes (Almeida, Moncada, & Bolaños, 2004; Herrero-Mendez et al., 2009; Sonntag et al., 2017). Astrocytes, despite being the most abundant cell-type in the central nervous system (Nedergaard, Ransom, & Goldman, 2003), have traditionally been considered support cells for neurons without significant relevance to neural function. However, recent work has determined critical roles of astrocytes in neural function including provisioning of metabolites to neurons for energy (Mächler et al., 2016; Pellerin & Magistretti, 1994; Volkenhoff et al., 2015) and enhancing synaptic processes (Diniz et al., 2012; Meyer-Franke, Kaplan, Pfieger, & Barres, 1995). These findings point to a need to characterize the important differences among a variety of cell types, not only in neurons, but in other metabolically-relevant brain cell types such as astrocytes between species to understand how the primate brain has evolved.

We hypothesize that there are important cell-type specific metabolic changes between human and chimpanzee brains and that astrocytes contribute, at least in part, to these differences. To investigate these changes, we used established protocols for the differentiation of induced pluripotent stem cells (iPSCs) into mature, functional neurons and astrocytes from humans and chimpanzees from multipotent neural progenitor cells (NPCs). This comparative cell culture approach allowed us to assess each cell type in the absence of other cell types in a defined, controlled environment. In order to determine adaptive interspecies differences in gene expression and metabolism in cell-type specific manner, we conducted RNA-Seq on human and chimpanzee NPCs, neurons, and astrocytes. We determined significant interspecies differential expression in all three cell types with the greatest degree of difference in astrocytes. Pathway enrichments revealed significant differences in cellular respiration between species across all cell types as well as cell-type specific changes in glucose and lactate transmembrane transport and pyruvate utilization suggestive of a higher capacity for energetic, rather than biosynthetic, metabolic phenotypes in human astrocytes. This work demonstrates a putative cell-type specific mechanism by which astrocytes may contributed substantially to the adaptive metabolic capacity of the human brain. It also contributes to a growing number of studies demonstrating the importance of considering astrocytes in presumably human-specific phenotypes, including neurodegenerative diseases.

## RESULTS

### RNA-Seq of human and chimpanzee iPSC-derived neural cells

We took a comparative genomics approach to investigating interspecies differences in neural cell-type specific gene expression between humans and chimpanzees. Three cell lines per species, representing three individuals, were used. These cell lines were originally obtained as fibroblasts from minimally invasive skin biopsies, reprogrammed into iPSCs, and have been validated for their pluripotency and differentiation abilities (Blake et al., 2018; Burrows et al., 2016; Eres, Luo, Hsiao, Blake, & Gilad, 2019; Pavlovic, Blake, Roux, Chavarria, & Gilad, 2018; I. G. Romero et al., 2015; Ward & Gilad, 2019; Ward et al., 2018). IPSCs from both species were initially cultured in the defined, iPSC-specific media mTeSR1 (STEMCELL, Vancouver, Canada). In order to investigate interspecies differences in cell-type specific gene expression between humans and chimpanzees, we generated RNA-Seq data from human and chimpanzee neural progenitor cells (NPCs), neurons, and astrocytes from induced pluripotent stem cells (iPSCs) (Figure 1A).

**Figure 1.**
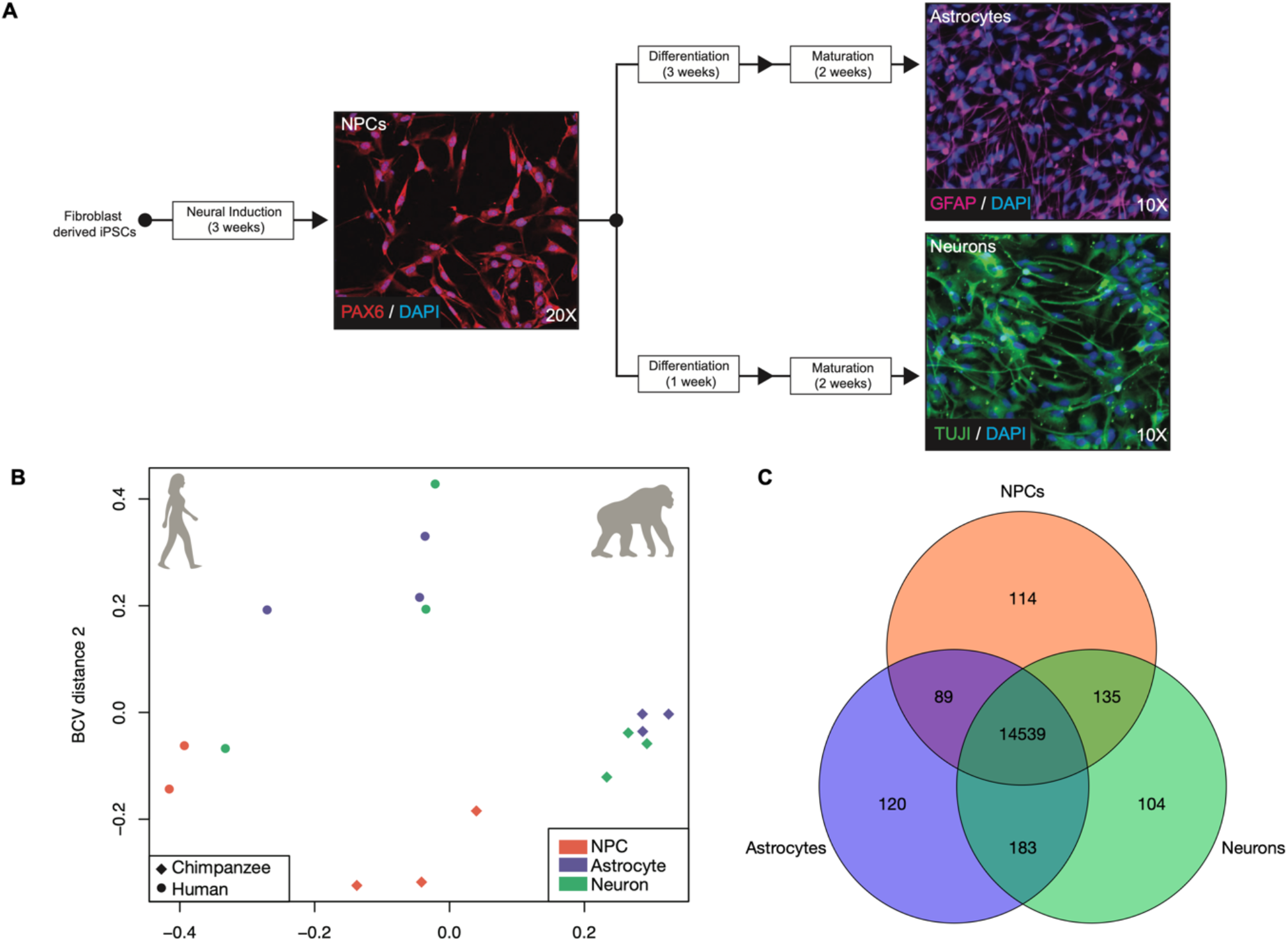
Patterns of gene expression variation of iPSC-derived neural cells from humans and chimpanzees. A) The differentiation schematic and representative immunofluorescent photos of iPSC-derived neural progenitor cells (NPCs; line H28815) stained with PAX6, astrocytes (line C3649K) stained with GFAP, and neurons (line C4955) stained with TUJ1. B) A PCoA of the iPSC-derived NPCs, neurons, and astrocytes transcriptomes. C) A Venn diagram of the overlap in expression across cell types. Further details for samples are included in SI Table 1.

To confirm that expression profiles of our iPSC-derived neural samples resembled that of neural tissue and primary neural cells, we used previously published data from human and chimpanzee tissues, including brain (Brawand et al., 2011), as well as human primary neurons and astrocytes (Materials & Methods) (Zhang et al., 2016). We visualized these data in an MDS plot and observed that our iPSC-derived neural samples clustered together within the same dimensional space as the other neural tissue and cell samples and not the non-neuronal tissue samples (SI Figure 5). These clustering analyses, in addition to *in vitro* validation of cell type by cell-type marker IHC prior to sequencing, demonstrate that we successfully created and obtained total transcriptome data of iPSC-derived NPCs, neurons, and astrocytes from humans and chimpanzees relevant for comparative assessments of cell-type specific interspecies gene expression differences.

### Astrocytes exhibit the greatest degree of differential expression between human and chimpanzee neural cell types

We next performed differential expression (DE) analyses in order to determine significantly differentially expressed genes between species. However, given the lack of clear distinction among our different cell types (Figure 1B), we first wanted to determine the degree of shared expression across all cell types. To do so, we determined overlap among cell types for genes with at least one count in one or more cell lines per cell type (Figure 1C). Of the total genes expressed in NPC (n=14,877), neuron (n=14,961), and astrocyte (n=14,931) samples, 95.13% (n=14,536) were shared among all three cell types (Figure 1C). This is consistent with previous findings that relatively few genes are cell-type specific in the brain, in terms of absolute expression (Magistretti & Allaman, 2015; McKenzie et al., 2018).

For DE analyses, we first conducted an analysis of variance (ANOVA)-like test for differentially expressed genes in a species (SP) by cell-type (CT) manner using edgeR (Robinson, McCarthy, & Smyth, 2010). We reasoned that this would be the most evolutionarily relevant set of genes for investigating neural cell-type specific “trade-offs” in expression between species. Using edgeR’s generalized linear model (GLM) functionality and a quasi-likelihood F-test for significant differential expression, we found 4,007 significantly differentially expressed genes in a species by cell type manner (26.22% of all expressed genes). However, at present, there are no post-hoc tests for an ANOVA-like test for differential expression, and so this analysis is limited in that it cannot delineate which samples (cell types) these genes are significantly DE in (Robinson et al., 2010). The ANOVA-like test for differences also requires an initial filtering of lowly expressed genes across all samples, which eliminates the 104-120 genes (0.68-0.79%, Figure 1C) expressed only in one cell type (CT-specific genes). For these reasons we also conducted interspecies pairwise DE comparisons for each CT (hereafter referred to as CT-DE analyses). While these CT-specific genes are relatively few in number, they likely have an important role in cellular function, and thus we did not want to exclude them from our interspecies CT-DE comparisons.

For CT-DE comparisons, the only genes included were those counts above zero in all samples per CT and were further filtered to those with counts per million (CPM) > 1 in at least 1 sample, resulting in 11,772 genes in NPCs, 12,451 genes in neurons, and 12,302 genes in astrocytes. We used the same GLM quasi-likelihood F-test to determine that 8.57% (n=1,294) of genes are differentially expressed between species’ NPCs, 5.8% (n=886) between neurons, and 12.2% (n=1,865) between astrocytes (Figure 2A, SI Figure 6). Many of these significantly differentially expressed genes in CT-DE comparisons overlapped with the SPxCT ANOVA-like differentially expressed genes (SI Table 3). When we determined overlap in differentially expressed genes between species across all three cell types, we found that, similar to global expression, a large number of genes were determined as differentially expressed between species in all three cell types (n=594, Figure 2B). However, there are far more genes that uniquely differentiate astrocyte gene expression between species (n=924) than NPCs (n=395) and neurons (n=100) (Figure 2B). This suggests that neuronal gene expression is more conserved across species in NPCs and neurons, and that astrocytes do indeed contribute to important interspecies differences in neural gene expression.

**Figure 2.**
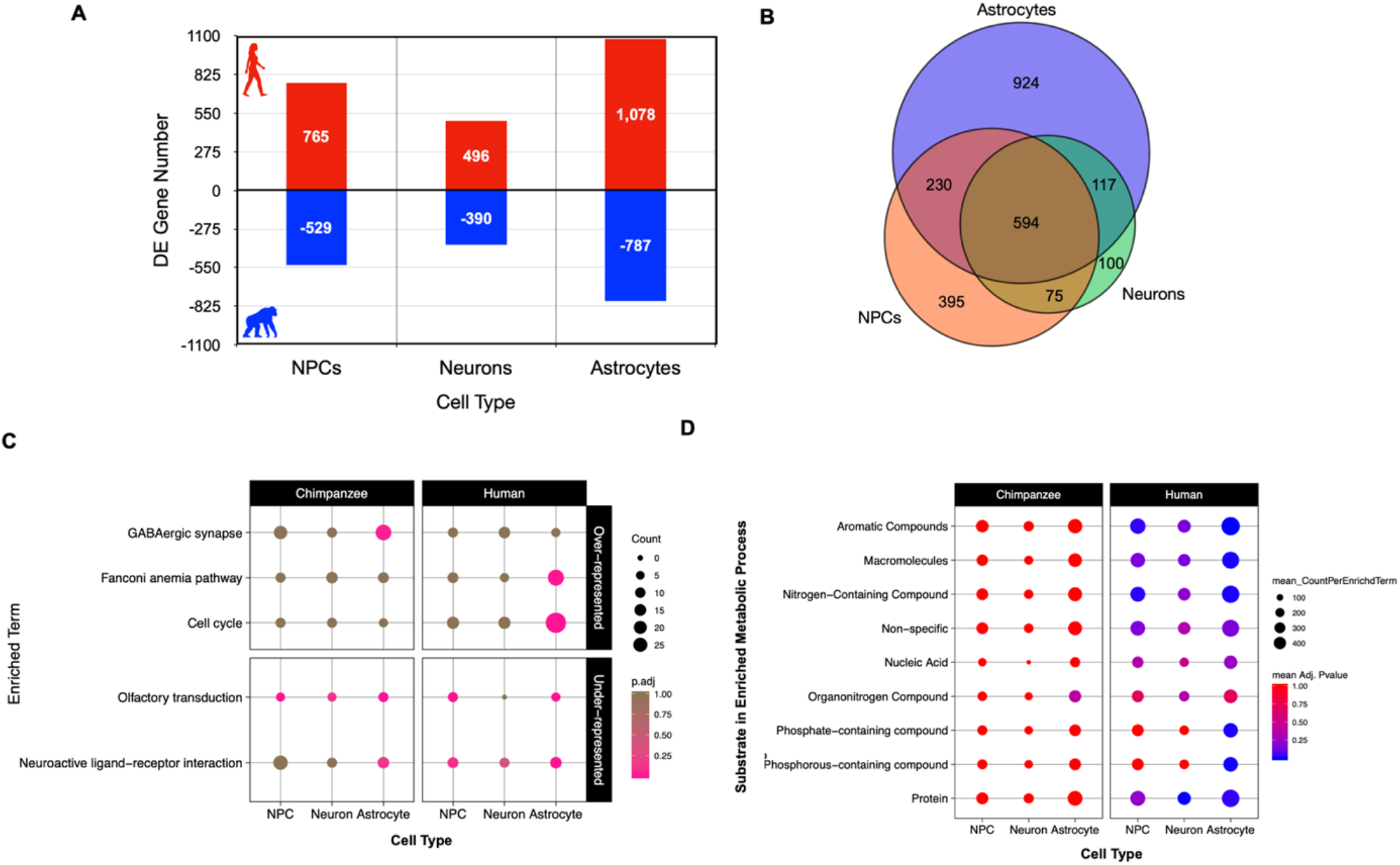
Astrocytes demonstrate the most significant differences in gene expression between human and chimpanzee neural cell types for metabolic but not neuron-specific pathways. A) Counts of genes exhibiting differential expression at an FDR < 5% between species for each cell type and the direction of higher expression for each CT-DE comparison (red/positive – higher expression in human; blue/negative – higher expression in chimpanzee). B) A Venn diagram of overlap in genes per cell type exhibiting differential expression between species. C) Plot of significantly (EASE < .05) over-represented (top panel) and under-represented (bottom panel) KEGG pathways determined by categorical enrichment analyses (full results in SI Table 4). Size indicates the count of genes per pathway while color indicates the adjusted p-value (pink – lower/significant, brown – higher/non-significant). D) Plot of significantly over-represented categories of GO BP terms determined by categorical enrichment analyses. (full results in SI Table 4). The categories (y-axis) represent groupings of multiple GO BP terms related to the metabolism of the indicated substrates/macromolecules. Size indicates the mean count and color indicates the mean adjusted enrichment p-value for all terms in that category. Refer to SI Table 4 and Methods for individual GO BP terms included in each category.

### Interspecies differences in gene expression are largely due to differential metabolic signaling skewed toward higher expression in humans regardless of cell type

We then used categorical enrichment analyses to determine what biological processes are over-represented (enriched) or under-represented (“conserved”) in interspecies differentially expressed genes by cell type (CT) (SI Table 4). There were consistently a larger number of interspecies differentially expressed genes per CT-DE comparison with higher expression in human cells (765 in NPCs, 496 in neurons, and 1,078 in astrocytes) than chimpanzee cells (529 in NPCs, 390 in neurons, and 787 in astrocytes) (Figure 2A). We used these six higher-in-one-species split DE gene lists in a mutliquery categorical enrichment analyses for under- and over-represented processes using gProfiler’s categorical enrichment tool (gOST) (Raudvere et al., 2019). Likely in part due to the larger number of genes with higher expression in human for all CT’s, there was consistently far more processes enriched in human CTs than chimpanzee CTs (SI Table 4).

Human and chimpanzee neural cells exhibited significant under-representation of the KEGG pathways ‘olfactory transduction’ and ‘neuroactive ligand-receptor interaction’ (Figure 2C). Consistently, both species’ also showed significant under-representation of GO biological processes (BP) terms related to development, immune function, and intracellular signaling (SI Figure 8). Human cells were under-represented for some extracellular and membrane associated cellular components (CC) (SI Figure 9) as well as molecular functions related to cytokine and receptor activity (primarily in astrocytes; SI Figure 10). Both species cells were underrepresented for nucleic acid binding and G-protein coupled receptor and transducer activity (SI Figure 10). This demonstrates that signaling, including some neuronal-specific signaling such as neuroactive ligand receptor interaction, and downstream perception processes (e.g. olfaction) are conserved across species for all cell-types.

As for significantly over-represented processes in differentially expressed genes between species, cell division, cytoskeletal and developmental signaling, and response to external stimuli terms were significantly over-represented in human astrocytes, transcription was enriched in human NPCs, and protein modification was enriched most significantly in human neurons (SI Figure 8). Because the human brain is so energetically demanding, we were specifically interested how pathways involved in cellular respiration and metabolism differed between species. Metabolic processes targeting a variety of substrates or macromolecules were enriched primarily in human cells (summarized in Figure 2D, full results in SI Table 4). Human astrocytes were significantly enriched for several more metabolic biological processes than human NPCs and neurons that included metabolism of phosphate-containing compounds as well as more generally for metabolism of macromolecules (Figure 2D). We also investigated enrichment of GO cellular component (CC) terms to determine if there were differences in expression of specific neuronal parts. Over-represented CC terms in human astrocytes were similar to the GO BP over-represented processes (cytoplasm, cytoskeleton, cell division and growth; SI Figure 9). Human neurons were enriched for terms related to intracellular macromolecule modification and trafficking (SI Figure 9B). Interestingly, human astrocytes were enriched for molecular functions (MFs) related generally to substrate binding, specifically, to ATP, carbohydrates and their derivatives, enzymes, and nucleic acids (SI Figure 10A). Human neurons were enriched for ubiquitin-related molecular activity, and human astrocytes for molecular activity related generally to ATPases, catalysis, exonucleases, helicases, kinases, and phosphotransferases (SI Figure 10B). These results indicate that metabolic processes differ between species in a CT-specific manner, and that all human neural cell types exhibit increased expression for a variety of macromolecular metabolic processes more than chimpanzee neural cells. Further, we see that there are significant differences in molecular functions important in cellular metabolic signaling.

### Human and chimpanzee neural cells differ in glucose and lactate transport as well as oxidative phosphorylation

Our results showed that when using unbiased categorical enrichment analyses without *a priori* expectations of enriched terms, metabolic processes targeting a variety of substrates or macromolecules were enriched in human neural cell types. However, very few of these processes were for pathways involved in cellular respiration resulting in production of energy in the form of ATP. There are known differences in metabolic capacity between neurons and astrocytes, including that astrocytes are characterized metabolically by high aerobic glycolytic activity (increased glycolysis with limited potential for oxidative ATP production) while neurons typically favor energy production and oxidative phosphorylation (reviewed in Magistretti & Allaman, 2015), we were interested in determining any interspecies, cell-type specific differences in these brain metabolic processes. To investigate if there were interspecies differences in expression of genes involved in aerobic glycolysis, we used a Gene Set Enrichment Analysis (GSEA) (Subramanian et al., 2005) with 23 *a priori* gene sets on the raw counts of the 12,407 genes used for interspecies pairwise CT-DE analyses. Gene sets were obtained from the Molecular Signatures Database (MSigDB) (Liberzon et al., 2011) and chosen in order to probe a variety of energetic metabolic pathways and substrate transporters of varying gene number size from multiple ontology categories (GO, KEGG, and REACTOME) (all probed gene sets listed in Figure 3A) (Antonazzo et al., 2017; Ashburner et al., 2000; Fabregat et al., 2018; Ogata et al., 1999). The goal was to determine if pathways involved in aerobic glycolysis (e.g. oxidative phosphorylation, glucose transport, TCA cycle) differ in a species by cell type manner, and so pathways not directly involved in aerobic glycolysis (e.g. fatty acid metabolism) are included as a comparison. We also included “control” pathways not directly related to metabolism (regulation of growth, and neurotrophin signaling).

**Figure 3.**
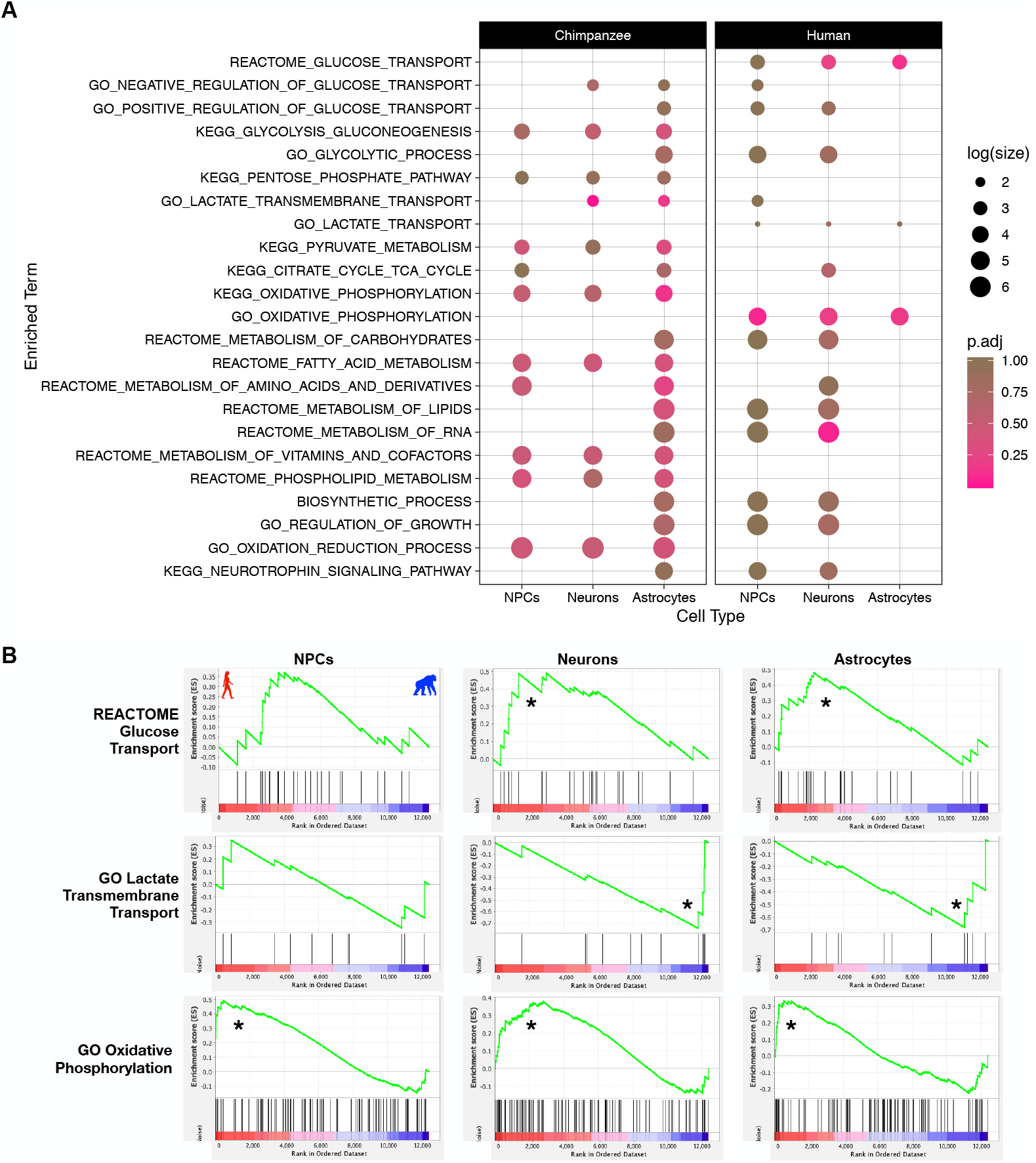
Humans and chimpanzees differ in metabolite transport and oxidative phosphorylation in a neural cell-type manner. A) Plot of all tested gene sets in the Gene Set Enrichment Analysis (GSEA) (full results in SI Table 5). Separate panels indicate which species ‘phenotype’ the gene set was enriched in. Color indicates the FDR Q-value (FDR < 25% indicates significance in this analysis). Size indicates the log(count) of genes included in the enriched gene set. B) Enrichment plots of significant pathways from GSEA for a subset of panel A. The green line indicates the running enrichment score for each gene in the gene set as the analysis moves down the ranked list of genes. The enrichment score for the gene set is the peak of this curve and an (*) indicates significantly enriched. The bottom panel is the ranked order of the genes and shows their location within that ranked set of genes. Left side of plot (and red/left portion of ranked order plot below) indicates human enrichment, while the opposite (right/blue) indicates chimpanzee enrichment. Full results in SI Table 5.

Our GSEA results indicate that the gene sets for lactate transmembrane transport, glucose transport, oxidative phosphorylation, and metabolism of RNA are significantly different between human and chimpanzee neural cells (FDR < 25% and nominal p-value < .05; Figure 3A, B, SI Table 5). Glucose transport was enriched in human neurons and astrocytes while lactate transmembrane transport was enriched in chimpanzee neurons and astrocytes (Figure 3A, B). The GO gene set for oxidative phosphorylation is significantly enriched in all human cell types while the KEGG oxidative phosphorylation is upregulated in chimpanzee astrocytes (Figure 3A, B). These results indicate cell-type by species differences in glucose uptake by cells, lactate shuttling, and diverging energetic cellular respiration.

### There are species by cell-type differences in expression of oxidative phosphorylation protein complexes

Leading edge analyses of significant GSEA gene sets are used to determine which genes of the gene set contribute most strongly to the enrichment of that pathway in the phenotype (Subramanian et al., 2005). We examined the results from the leading edge GSEA analysis with the CT-DE expression analyses to get a better idea of how these three pathways diverge in a cell type by species manner (Figures 4 and 5; full results in SI Table 6). We calculated a rank for DE genes for each CT-DE comparison (NPC, neuron, and astrocyte): (sign of logFC) x log10(FDR Q-value) (Reimand et al., 2019) and used that in addition to the GSEA leading edge analysis to determine significant differences. For the oxidative phosphorylation genes, we were interested in determining why there was a difference in species and cell type enrichment based on the source of the gene set (KEGG vs GO) and determining if there were potential functional differences in oxidative phosphorylation between species. We mapped the CT-DE rank of the core enriched genes for GO (Figure 4A) and KEGG (Figure 4B) oxidative phosphorylation genes.

**Figure 5.**
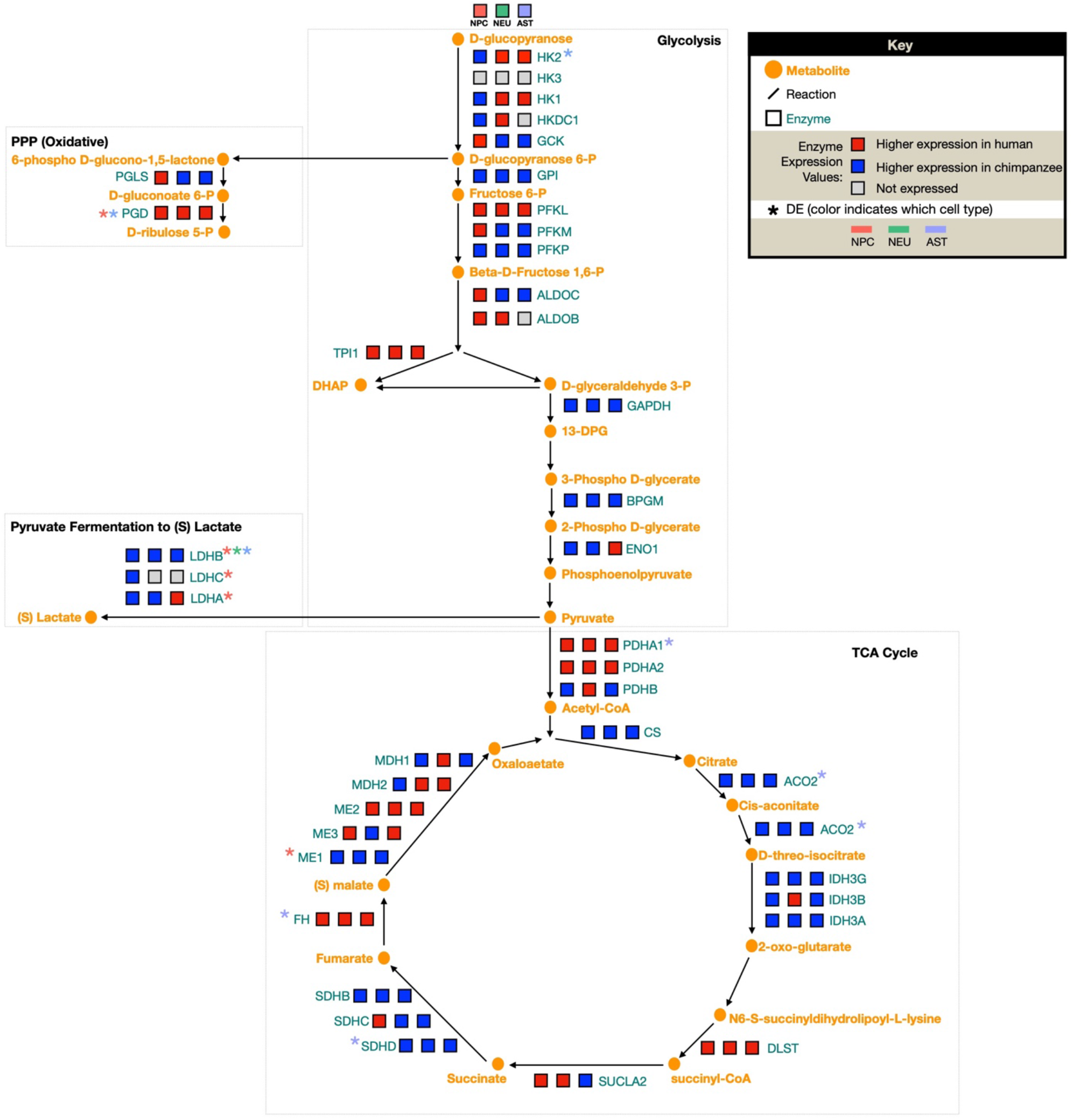
Divergence in pyruvate utilization between species’ astrocytes. We constructed a focal set of aerobic glycolysis signaling pathways in order to contextualize our DE results in the framework of a network signaling. A diagram of the major pathways involved in aerobic glycolysis (glycolysis, pentose phosphate pathway (PPP), lactate conversion from pyruvate, and TCA cycle). For each enzyme in the pathway, three blocks indicate expression of this enzyme in each cell type – left to right: NPCs, neurons, astrocytes. Color indicates level of expression (higher in human (red), higher in chimpanzee (blue), not expressed in this cell type (grey)).

The core set of genes in GO and KEGG oxidative phosphorylation (OXPHOS) gene sets included genes for subunits of cytochrome c oxidase (the nuclear-encoded *COX4I1, COX6B2, COX7B*, and *COX7C* and the mitochondrially-encoded *MT-CO1, MT-CO2*, and *MT-CO3*) as well as those that aid in cytochrome c oxidase assembly (*COX10* and *COX11*) (Figure 4A, B; SI Table 6). Cytochrome C oxidase is the terminal complex in the electron transport chain and is crucial to maintaining a proton gradient across the inner mitochondrial membrane for ATPase to synthesize ATP. These genes are of particular interest, because within anthropoids, genes encoding the subunits of cytochrome c oxidase show an accelerated rate of evolution in their sequences compared with any other placental mammals (Preuss, 2012). Here, we see a cell type by species divergence in cytochrome c oxidase gene expression, where most of these genes exhibit higher expression in human neurons (Figure 4A). There is also a clear trend of mitochondrially-encoded genes that function in OXPHOS having significantly higher expression in human cells, including those for cytochrome c oxidase subunits (*MT-CO1, MT-CO2, MT-CO3*), but also mitochondrially-encoded ATP synthase (*MT-ATP6*) and mitochondrially-encoded subunits of the NADH:ubiquinone oxidoreductase core of electron transport chain complex I (*MT-ND2, MT-ND3, MT-ND5*, and *MT-ND6*) (Figure 4A).

**Figure 4.**
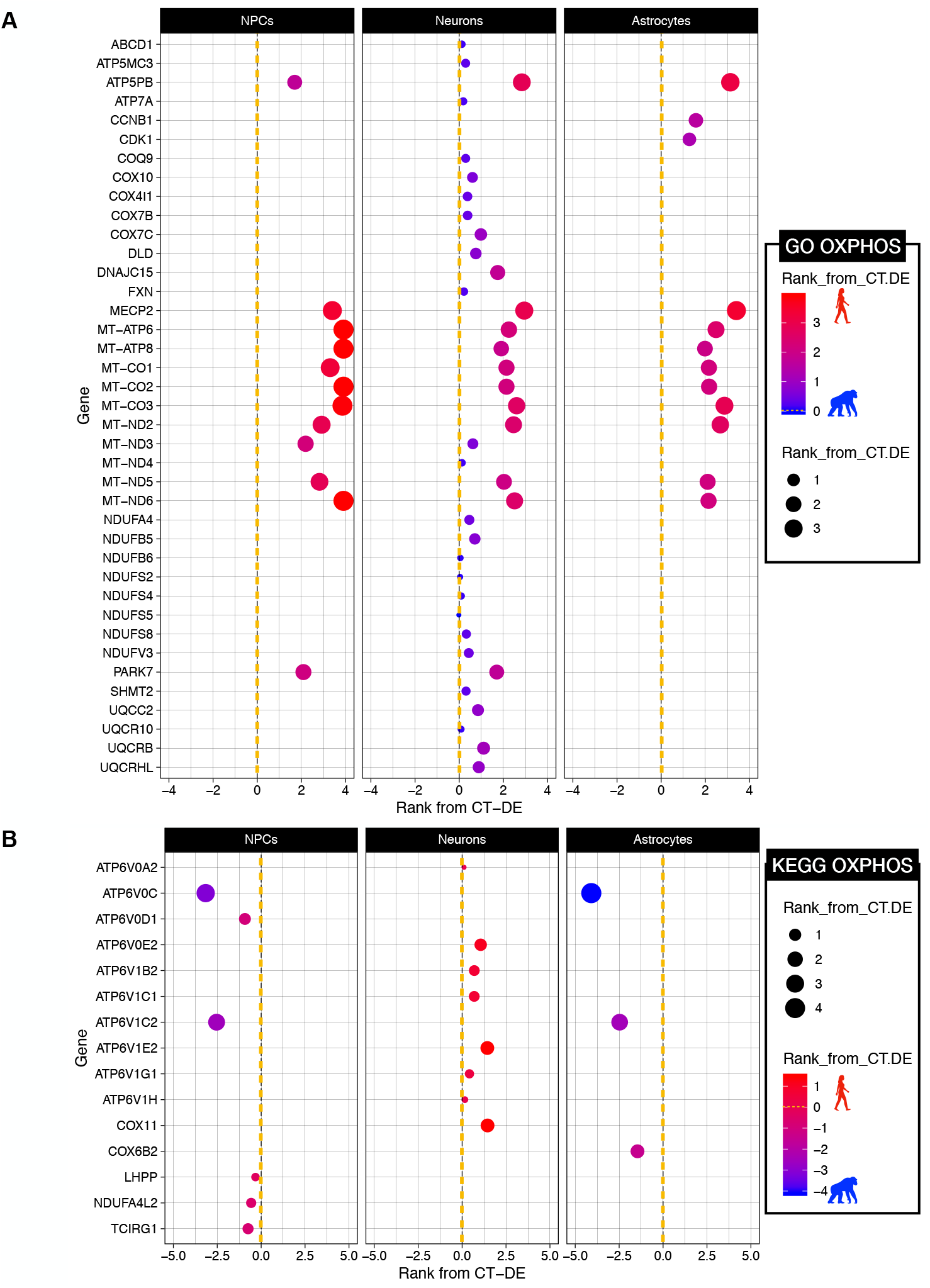
Interspecies expression differences of oxidative phosphorylation genes is influenced by higher expression of mitochondrial genes in all neural human cell types. CT-DE results of genes per A) GO or B) KEGG oxidative phosphorylation gene sets determined as members of the core set of genes influencing significant enrichment of these gene sets in the GSEA analysis. CT-DE rank was calculaterd per each gene [(sign of logFC) x log10(FDR Q-value)], with values greater than zero indicating higher expression in human and values less than zero indicate higher expression in chimpanzee. Color spectrum and size also indicate rank (red – higher in human, blue – higher in chimpanzee, larger = higher rank). See SI Table 3 for DE results per gene and SI Table 6 for GSEA results per gene.

A major difference between the GO and KEGG OXPHOS gene sets is that the KEGG OXPHOS set includes vacuolar-ATPase (V-ATPase) genes, whose major role is in acidification of intracellular organelles, and have an important function in synaptic vesicle proton gradient formation and maintenance (Maxson & Grinstein, 2014; Pamarthy, Kulshrestha, Katara, & Beaman, 2018). There is an intriguing pattern of enrichment for higher expression of subunits of V-ATPases in a cell type by species manner (Figure 4). Three genes for subunits of vacuolar ATPases (*ATP6V0C, ATP6V0D1*, and *ATP6V1C2*) are core enriched genes in the KEGG OXPHOS gene set and are significantly enriched in CT-DE with higher expression in chimpanzee NPCs and astrocytes, but not neurons (Figure 4B). Furthermore, only one of these vacuolar-ATPase genes is DE in a SPxCT manner (*ATP6V1C2*) (Figure 4B). However, several other V-ATPase subunit genes are core enriched only in human neurons (Figure 4B), most notably *ATPV1E2* and *ATPV0E2*, both of which are core enriched and significantly differentially expressed in human neurons. This shows that V-ATPases exhibit significant DE between humans and chimpanzees, and that human neurons are distinct in V-ATPase gene expression from chimpanzee NPCs and astrocytes. Given the important function of V-ATPases in synaptic vesicle formation for neurotransmitter signaling, this may imply an important functional change in humans specifically in neurons.

### Interspecies differential expression of important metabolite transporter genes in neurons and astrocytes

It is widely accepted that neurons exhibit limited glycolytic capacity and that astrocytes respond to signals associated with increased synaptic signaling by increasing glucose uptake and subsequent aerobic glycolysis of glucose to produce lactate to be used as energy source by neurons (reviewed in Magistretti & Allaman, 2015; Pellerin & Magistretti, 1994). For this reason, we were interested in investigating if there were interspecies gene expression differences in lactate transport, particularly in neurons and astrocytes. We performed GSEA using glucose and lactate transport pathways. The GO gene set ‘lactate transmembrane transport’ was enriched in chimpanzee neurons and astrocytes, showing that genes involved in lactate transport are more highly expressed in these mature cell types than NPCs (SI Figure 11A). Several genes in this gene set are for proton-linked monocarboxylate transporters that transport pyruvate and lactate (*SLC16A11, SLC16A12, SLC16A13*, and *SLC16A6*) that are all core enriched in neurons and astrocytes (SI Figure 11A). *SLC6A11* and *SLC16A13* are also differentially expressed in SPxCT ANOVA-like DE as well as an CT-DE manner, though *SLC16A13* is not in astrocytes (SI Figure 11A). The enrichment for the lactate transmembrane transport gene set in chimpanzee neurons and astrocytes and the corresponding DE of specific pyruvate and transporter genes between species’ may suggest that chimpanzee neurons and astrocytes have the capacity to shuttle pyruvate and lactate at a higher rate than human neurons and astrocytes.

In addition to lactate transmembrane transport enrichments, the glucose transport gene set was significantly enriched in human neurons and astrocytes (Figure 3). There were two hexokinase genes (*HK1* and *HK2*) core enriched in this gene set that demonstrate lower expression in chimpanzee NPCs but higher expression in human astrocytes (SI Figure 11B), though only *HK2* is significantly upregulated in human astrocytes by CT-DE analysis (SI Figure 11B). *G6PC3* may not be significantly differentially expressed in any particular cell type, but it is in the SPxCT DE comparison and does show insignificant but consistently higher expression in all chimpanzee cell types (SI Figure 11B). *SLC2A3* is a facilitative glucose transporter across the cell membrane, and here, it exhibits core enrichment in all three human cell types by the GSEA leading edge analysis, as well as moderately (though non-significant) higher expression in human (SI Figure 11B). The enrichments of glucose transport in human neurons and astrocytes appears to be influenced by increased expression of plasma membrane associated glucose transporters (e.g. *SLC2A3*) and enzymes that function in the earlier steps of glycolysis (*HK1, HK2, G6PC3*).

### A subset of genes exhibiting significantly higher expression in human astrocytes also have signs of positive selection in their promoter regions

In order to begin to probe whether expression differences between species are influenced by selective pressures, we obtained synonymous (dS) and nonsynonymous (dN) nucleotide mutation rates from Ensembl (Kersey et al., 2017; Schneider et al., 2017) and compared the rate of change (dN/dS) for different groups of iPSC-derived neural cell expressed genes. A dN/dS > 1 indicates putative evidence of positive selection in coding regions (Herrero et al., 2016). As predicted, the vast majority of all the genes identified as expressed in these cells did not exhibit a dN/dS > 1 (SI Figure 12). Only few genes DE between species in iPSC-derived astrocytes (n=6), neurons (n=6), and NPCs (n=11) exhibit signs of coding selection (dN/dS > 1) (SI Table 7, SI Figure 12). The gene *HRC* (histidine rich calcium binding protein) exhibits positive selection and is significantly DE between species in NPCs, but not astrocytes, and is not expressed at all in neurons (SI Table 7). Three genes (*DCTN6, HHLA3, DBNDD2*) have a dN/dS > 1and showed significantly DE in all three cell types (SI Table 7). However, there is no commonality in these genes to suggest any meaningful impact on gene expression differences or in specific cell types. This is likely due, in part, to the limitations of the methods used for probing coding sequences, rather than those in non-coding, *cis*-regulatory changes, which previous studies have demonstrated are critical for significant differences in expression in primate brains (Babbitt et al., 2010; Bauernfeind et al., 2015; Haygood et al., 2007).

To further probe for signs of selective pressure in non-coding regions, we tested the promoter regions of aerobic glycolysis genes that were expressed in our samples. Thirteen of 156 aerobic glycolysis genes exhibited signs of positive selection (SI Table 8). These included two V-ATPase component proteins (*ATP6V1G1* and *ATP6V1H*), four nucleoporins (*NUP85, NUP54, NUP214*, and *NUP107*), a subunit of the NADH dehydrogenase complex of the ETC (*NDUFA4*), cyclin B1 (*CCNB1*), an RNA binding protein (*RAE1*), and two glycolysis genes, glucokinase regulator (*GCKR*) and hexokinase (*HK1*). Interestingly, though all of these genes were expressed to some degree in all three cell types and in both species (with the exception of *GCKR*), they were only ever significantly differentially expressed between species in astrocytes (n=4 DE in astrocytes; SI Table 8). Of these four genes that were significantly DE between species and under positive selection include *CCNB1, NDUFA4*, and *NUP85* were more highly expressed in human astrocytes, while *SLC16A11* was more highly expression in chimpanzee astrocytes. Of note, *GCKR* is only expressed in astrocytes, with significantly higher expression in chimpanzee (SI Table 8). Significant results from this test suggest regulatory elements that control expression of these genes may be under selection in humans. Significant results from this analysis support selection in genes involved in metabolic processes in humans.

### Differences in the aerobic glycolysis genes are primarily in NPCs and astrocytes but not neurons

In order to obtain a more comprehensive, pathway-level understanding of altered expression of aerobic glycolysis in iPSC-derived neural cells between humans and chimpanzees, we reconstructed a signaling network diagram of enzymes involved in four sub-pathways involved in aerobic glycolysis (glycolysis, pentose phosphate pathway, pyruvate conversion to lactate, and the citric acid (TCA) cycle) from the HumanCyC database (P. Romero et al., 2005) (Figure 5). We then mapped discrete expression values (higher in human, higher in chimpanzee, not expressed) for each of these enzymes in all three cell types onto the pathway diagram to illustrate which species the enzymes were more highly expressed in and if they were significantly differentially expressed between species. From this, we see dynamic changes in expression across aerobic glycolysis sub-pathways, with no significant shift towards higher expression of enzymes in one species or cell type at any of these sub-pathways (Figure 5). Aerobic glycolysis enzymes exhibiting interspecies DE in NPCs were *PGD, LDHA, LDHB, LDHC*, and *ME1*, enzymes demonstrating DE in astrocytes were *PGD, HK2, PDHA1, FH, ACO2*, and *SDHD*, and only a single enzyme exhibited interspecies DE in neurons (*LDHB*) (Figure 5). This shows that NPCs and astrocytes, but not neurons, exhibit the vast majority of significant differences in expression of enzymes in these pathways (Figure 5). The majority of these genes were expressed in all cell types, particularly those that exhibited significant DE in NPCs or astrocytes, so this lack of DE in neurons is not simply due to cell-type specific expression differences (SI Figure 13). Human and chimpanzee astrocytes appear to diverge at the stage of pyruvate utilization, where human astrocytes exhibit significantly higher expression of *PDHA1*, which converts pyruvate into acetyl-coA whereas chimpanzee astrocytes show significantly higher expression of *LDHB*, which converts pyruvate into lactate rather than acetyl-coA (Figure 5). Interestingly, *LDHB* is also the only enzyme in these pathways exhibiting differential expression between species in neurons. Other *LDH* isoforms (*LDHC* and *LDHA*) also exhibit significant interspecies DE, with higher expression in chimpanzee NPCs. All chimpanzee neural cell types differ from human chimpanzees for *LDH* expression, but chimpanzee NPCs differ from human NPCs in expression levels of multiple *LDH* isoforms. This pathway level consideration of expression differences between species suggests significant changes in aerobic glycolysis enzyme activity primarily in NPCs and astrocytes and an interspecies divergence in pyruvate utilization.

## DISCUSSION

Our novel approach using iPSCs allowed us to investigate rare neural cell types from primates to determine cell type specific metabolic changes necessary to support evolution of the human brain. Our results demonstrate that interspecies divergence in gene expression is more conserved in neurons and significantly greater between species’ astrocytes. Differential expression between species’ cell types is enriched for metabolic processes related to cellular respiration. This finding is similar to that of previous studies of differential expression between human and chimpanzee whole brain tissue (Haygood et al., 2007; Kosiol et al., 2008; Uddin et al., 2008), and our results suggest this is driven primarily by higher expression of metabolic genes in human cells. However, there were some interesting examples of potential tradeoffs in expression patterns of specific genes and pathways in a cell-type by species manner. We determined that human neurons and astrocytes are enriched for higher expression of glucose transport proteins while chimpanzee neurons and astrocytes exhibit higher expression of lactate transmembrane transport genes and that there are dynamic interspecies changes in expression of nuclear- and mitochondrially-encoded subunits of the protein complexes important for oxidative phosphorylation. Our study demonstrates the utility of iPSC-derived cells for better understanding evolution of gene expression in primate brains.

Previous work has determined several significant differences in expression of cellular respiration pathways and evidence of differential selective pressure associated with metabolic genes (both noncoding and coding) between human and chimpanzee brains (Goldberg et al., 2003; Grossman et al., 2001; Grossman et al., 2004; Hüttemann et al., 2012; Uddin et al., 2008; Wildman et al., 2002; Wu et al., 1997). However, the heterogenous nature of brain tissue has complicated drawing conclusions about the contribution of specific cell types to this trajectory of elevated metabolic expression in human brains. Specifically, there is a long-standing question about the sole influence of greater neuron numbers in human brains (Herculano-Houzel, 2011) on the observed increase in glucose utilization (Hofman, 1983; Mink et al., 1981). Our approach allowed us to further investigate the cell-type specific contributions to long-understood differences in brain metabolic capacity. Because our methods of determining signficant differences in gene expression do not rely on number of cells or absolute quantity of transcripts, we are able to conclude that there are cell-type specific contributions to altered metabolic gene expression between species’ neural cell types, and that sheer number of cells alone likely does not fully explain metabolic differences between human and chimpanzee brains. We found that astrocytes display the greatest proportion of interspecies difference in metabolic gene expression and that neuronal gene expression actually appears to be more conserved across species. Our results demonstrate that, in light of the relatively recent discovery of cell-type specific metabolic differences between neurons and astrocytes, investigation of differences in brain metabolism among primates and the evolutionary processes that shaped them would indeed be incomplete without the consideration of all neural cell types, not just neurons. This suggests that astrocyte-mediated differences in metabolic brain function may be an important mechanism by which the ultimate evolutionary trajectory of human brain evolution has occurred.

Human cells show increased capacity for glucose transport, via greater expression of glucose transporters, in the mature neural cell types investigated here. This may imply that the observed differences in glucose utilization of the human brain extend beyond development and may play an important role in more mature neurons and astrocytes for either energy or macromolecule production. Furthermore, lactose dehydrogenase (*LDH*) isoforms favor differential affinities for interconverting pyruvate and lactate. *LDHB* favors the production of lactate into pyruvate (Almad et al., 2016; Bauernfeind & Babbitt, 2014), and is significantly differentially expressed with higher expression in chimpanzee for all cell types. This coupled with higher expression of lactate transporters in chimpanzee astrocytes suggests that chimpanzee cells may be favoring production and transport of lactate at a higher rate than all human neural cell types tested. This opposing enrichment for elevated glucose transport in mature human neural cell types in comparison to elevated lactate transport and conversion to pyruvate in mature chimpanzee cells raises some intriguing questions about metabolic trade-offs between human and chimpanzee brains. If we presume that the direction of change in metabolic gene expression is on the human lineage, and we do have some evidence from signs of positive selection on glucose and energetic metabolism coding and non-coding genes within primates with proximity to humans (Goldberg et al., 2003; Grossman et al., 2001; Grossman et al., 2004; Hüttemann et al., 2012; Uddin et al., 2008), then perhaps an increase in glucose uptake in human brains has allowed for a decrease in expression of genes that convert and shuttle lactate (via LDH and lactate transporters) to produce pyruvate. Previous studies have found lineage-dependent differences in enrichments for metabolic pathways in genes DE between primate brain regions, with greater glucose and carbohydrate metabolism in humans but higher glycogen and acyl-CoA metabolism in chimpanzees (Haygood et al., 2007; Kosiol et al., 2008; Uddin et al., 2008). The increase in LDH and lactate transport in chimpanzee neurons and astrocytes may be an important cell-type specific mechanism contributing to findings of significant metabolic differences in previous studies of whole brain tissue.

Human neural cells were enriched at the pathway level for oxidative phosphorylation genes, and within that pathway, there were some interesting examples of opposing enrichment for subunits of oxidative phosphorylation protein complexes. We observed increased expression and enrichment for components of cytochrome c oxidase, which previous studies have determined genes involved in this complex to be under positive selection (Goldberg et al., 2003). However, we expand on the knowledge of interspecies differences in cellular respiration complex expression by determining that these components are more highly expressed in all human cell types investigated, and particularly in human neurons (Figure 4A). We also observed higher expression for subunits of other electron transport chain complexes, including ATP-synthase and the NADH:ubiquinone oxidoreductase components of complex I. This cell-type by species approach also allowed for us to determine that human neurons and chimpanzee NPCs and astrocytes have higher expression for genes involved in vacuolar-ATPase function. We also determined that two V-ATPase genes exhibit signs of positive selection in their coding regions, though neither were differentially expressed between species. Our findings that V-ATPases are significantly differentially expressed between humans and chimpanzees suggests that human neurons are distinct in V-ATPase gene expression from chimpanzee NPCs and astrocytes. Given the important function of V-ATPases in synaptic vesicle formation for neurotransmitter signaling, this may be an mechanism by which human-specific changes in neuronal signaling has occurred.

Our investigation into the overlap of signatures of positive selection in coding regions of genes exhibiting interspecies DE revealed very little new or intriguing information. This is likely due at least in part to the limited scope of the current methods for searching for positive selection. The dN and dS scores obtained from Ensembl for use in this analysis were averages across all sites in a given gene, thus minimizing significant changes at specific sites (Yang, 2007). Our investigation of non-coding regulatory regions of genes in oxidative phosphorylation and glycolysis pathways found several genes to be under positive selection to varying degrees in humans, with some overlap with previous reports (Haygood et al., 2010) though there were some noticeable differences. For example, glucose-6-phosphate isomerase (*GPI*), which has been determined to be under positive selection in its non-coding promoter region in previous studies (Haygood et al., 2007), did not exhibit interspecies DE in any of these cell types. However, we did find evidence of positive selection in promotors of several aerobic glycolysis genes in the human lineage. Interestingly, we see that aerobic glycolysis genes exhibiting positive selection in promoter sequences were only significantly differentially expressed between species in astrocytes, not NPCs or neurons. Hexokinase and glucokinase both function in the conversion of glucose to glucose-6-phosphate, thus playing an early role in glycolysis. We found that there are significant expression differences in these key genes between species’ astrocytes. Human astrocytes significantly upregulate *HK1* while chimpanzee astrocytes significantly upregulate glucokinase regulator (*GCKR)*, which also exhibits positive selection in its promoter. This suggests that there is an evolved difference in the initial processing of glucose during glycolysis in an astrocyte-mediated manner, in addition to interspecies differences in the expression of glucose transporters. We also found evidence of adaptive divergent astrocyte glycolytic activity between species’ for utilizing pyruvate. In addition to the difference in pyruvate conversion enzymes and lactate transmembrane shuttling, we found evidence of positive selection on the human lineage and a significant increase in expression in chimpanzee astrocytes of *SLC16A11*, which functions in catalyzing transport of pyruvate across the plasma membrane. These results are intriguing in that they demonstrate that evidence of positive selection in the human lineage and divergent gene expression in genes involved in pyruvate processing and transport. These positive selection analyses further corroborate that there are significant differences in glycolytic gene expression between species’ astrocytes at initial steps in glycolysis as well as pyruvate utilization. Combined, this supports the evolution of metabolism in the human brain. Future investigations of the overlap between genes exhibiting DE in a cell-type specific manner and signatures of positive selection should utilize methods that allow for branch and site models that are more effective at determining positive selection in a lineage-specific manner (e.g HyPhy for non-coding and coding regions) (Haygood et al., 2007; Horvath et al., 2014; Muntané et al., 2014; Pond, Frost, & Muse, 2004).

Our focal analysis of aerobic glycolysis enzyme expression yielded several important findings. We show that there is not a consistent single species skew in expression levels for any of the sub-pathways in aerobic glycolysis (e.g. all glycolysis enzymes exhibited higher expression in one species or another, with no change in genes involved in TCA or lactate shuttling). The lack of significant DE between species in neurons for aerobic glycolysis enzymes demonstrates the importance of studying cell types other than neurons when investigating human brain evolution, and suggests that astrocytes may indeed be critical for the evolution of the metabolically demanding human brain. We also found that it is not simply the lack of glycolytic capacity of neurons (Almeida et al., 2004; Herrero-Mendez et al., 2009; Sonntag et al., 2017) that contributes to cell-type specific signaling disparities, at least in a comparative manner, because the majority of interspecies differentially expressed genes were expressed to some degree in all cell types.

Perhaps the most intriguing finding is the interspecies divergence in processing pyruvate. Humans exhibit significantly higher expression of *PDHA1* than chimpanzees do, indicative of a functionally relevant increase in conversion of pyruvate into acetyl-CoA and further utilization of the products of glycolysis for energy production, while chimpanzee astrocytes exhibit expression phenotypes suggestive of greater lactate production (higher expression of *LDH*, which converts pyruvate to lactate) as well as enrichment for greater lactate transmembrane transport in chimpanzee neural cell types (higher expression of lactate transmembrane transporters). This suggests that chimpanzee neural cells, and most prominently astrocytes, have a significantly greater capacity to convert pyruvate into lactate and then shuttle it across membranes than human astrocytes do. These analyses suggest significant interspecies changes in aerobic glycolysis enzyme activity primarily in NPCs and astrocytes and an interspecies divergence in pyruvate utilization. Previous work has shown a shift from aerobic glycolysis in NPCs to oxidative phosphorylation in more mature neurons (Zheng et al., 2016), but this study is the first of our knowledge to compare across species and include astrocytes. More generally, we see that astrocytes exhibit the greatest degree of expression difference between species than the other cell types while neuronal gene expression is more conserved. A recent investigation of multiple brain regions from human, chimpanzee, bonobo, and macaque using single-cell RNA-seq also found that astrocytes were one of the cell types exhibiting the greatest expression differences in humans (Khrameeva et al., 2020). This increased variation in interspecies gene expression in astrocytes suggests that previously observed differences in whole brain gene expression may be due astrocyte-specific changes to a larger degree than previously thought, and that this is a crucial cell type to consider when investigating human-specific brain gene expression has evolved.

Evolved differences in metabolic investment may be the basis for a number of primate-specific phenotypes, including those that are unique to humans, (e.g. slow reproduction and long lifespan (Charnov & Berrigan, 1993; Pontzer et al., 2014; Snodgrass, Leonard, & Robertson, 2007)). Our results provide insight into the metabolic changes that were necessary to support evolution of the human brain. We have demonstrated a significant interspecies divergence in aerobic glycolytic gene expression in astrocytes, suggesting that this traditionally understudied glial cell type likely contributes to the tissue-level shifts in gene expression and that astrocytes play an important role in the evolution of the metabolically expensive human brain. A potential challenge in cell-type specific studies of interspecies differences in brain gene expression is the loss of intercellular signaling between different cell types, a hallmark of synaptic signaling in whole tissue. Furthermore, the astrocyte-neuron lactate shuttle links the complementary metabolic needs of astrocytes and neurons (reviewed in Magistretti & Allaman, 2015; Pellerin & Magistretti, 1994). Future studies of gene expression differences with controlled levels of intercellular signaling by building in complexity (e.g. interspecies differences in expression of single cell types compared to that of co-cultured iPSC-derived neurons and astrocytes) could further inform interspecies differences in neuronal gene expression.

## MATERIALS AND METHODS

### SAMPLES AND CELL CULTURE

Induced pluripotent stem cells (iPSCs) from three individuals (cell lines) per species (human and chimpanzee) were cultured in defined, iPSC-specific media mTeSR1 (STEMCELL, Vancouver, Canada). These cell lines were originally obtained as fibroblasts from minimally invasive skin biopsies, reprogrammed into iPSCs, and have been extensively validated for their pluripotency and differentiation abilities (Blake et al., 2018; Burrows et al., 2016; Eres et al., 2019; Pavlovic et al., 2018; I. G. Romero et al., 2015; Ward & Gilad, 2019; Ward et al., 2018). Three cell lines per species, representing three male individuals, were used (SI Table 1). To investigate differences between human and chimpanzee neural cell types, we induced iPSCs from each species first into multipotent, neural-lineage committed neural progenitor cells (NPCs) using STEMdiff Neural Induction Medium in monolayer for three passages (21-28 days), as per manufacturer’s instructions (STEMCELL Technologies, Vancouver, Canada). Successful transition of iPSCs into NPCs was determined using immunofluorescence for the absence of the stem-cell marker OCT4 and presence of the NPC-marker PAX6 (Figure 1A). NPCs were then expanded into three subsets: one for RNA collection, and two for further differentiation and maturation into neurons and astrocytes. We then differentiated NPCs into mature neurons and astrocytes using the neuron and astrocyte specific STEMdiff differentiation and maturation kits as recommend by the manufacturer. Briefly, we differentiated NPCs using the STEMdiff Neuron Differentiation Medium for one week and then matured them using the STEMdiff Neuron Maturation Medium for two weeks. Similarly, we differentiated NPCs using the STEMdiff Astrocyte Differentiation Medium for three weeks and then matured them using the STEMdiff Astrocyte Maturation Medium for two weeks. All cells were validated for cell type via immunofluorescence prior to harvesting as follows: NPCs for PAX6+/OCT4-(Developmental Studies Hybridoma Bank, Iowa City, IA), neurons for neuron-specific class III β-tubulin (TUJ1; Neuromics, Edina, MN), and astrocytes for GFAP (Sigma Aldrich, St. Louis, MO), according to manufacturer’s suggestions (SI Figure 1). All mature iPSC-derived cells for each cell type were harvested at similar timepoints: NPCs at passage 5-6 post-induction from iPSCs, mature neurons at passage 3-4 and mature astrocytes at passage 5-6 post-differentiation from NPCs and subsequent maturation (SI Table 1). We used edgeR (Robinson et al., 2010) to normalize our raw counts across all samples and visualized these data using a multidimensional scaling (MDS) plot of all of the expressed genes (Figure 1B).

### LIBRARY PREPARATION AND SEQUENCING

Total RNA was extracted from cells (1-2 wells, 6 well plate) using an RNeasy Plus Mini Kit (Qiagen, Hilden, Germany), including a DNase step to remove residual DNA. Total RNA was analyzed for quality using the Bioanalyzer RNA 6000 Nano kit (Agilent, Santa Clara, CA) with RNA Integrity Numbers (RINs) for all samples between 8.3-10 (SI Table 1). Using the NEBNext Poly(A) Magnetic mRNA Isolation Kit (NEB), mRNA was isolated from intact total RNA, and cDNA libraries were made from each sample using the NEBNext RNA Ultra II Library Prep Kit for Illumina (New England Biolabs, Ipswich, MA). Barcoded samples were sequenced using the Illumina NextSeq 500 (Illumina, San Diego, CA) platform at the Genomics Resource Core Facility (Institute for Applied Life Sciences, UMass Amherst) to produce 75 base pair single-end reads, yielding a minimum of 32 million reads per sample.

### READ MAPPING AND QUANTIFICATION

Quality-filtered reads were aligned to respective species’ most recent ENSEMBL genome (*Homo sapiens* GRCh38 and *Pan troglodytes* PanTro3.0 (Kersey et al., 2017; Schneider et al., 2017)) with Bowtie2 (Langmead & Salzberg, 2012) using default ‘--local’ parameters for gapped alignments, with a minimum alignment percentage of ≥ 98.84% (SI Table 1). HT-Seq (Anders, Pyl, & Huber, 2015) was used to quantify counts per gene for each sample, using ENSEMBL gene transfer files (GTFs) corresponding to the same genome build used for alignment (Aken et al., 2016). High quality, one-to-one orthologs from *P. troglodytes* were matched to the ENSEMBL human reference set of genes using biomaRt (Kinsella et al., 2011), yielding 15,284 genes identified as expressed in at least one sample. All data are available in FASTQ format in the National Center for Biotechnology Information’s (NCBI) Short Read Archive (SRA) with accession number PRJNA665853 (publicly available upon publication). A link for reviewers’ pre-publication can be found here: https://dataview.ncbi.nlm.nih.gov/object/PRJNA665853?reviewer=hobvb5i4ejtpt0jp6r9r29e372.

### CLUSTERING ANALYSES

We used clustering analyses to determine the variation among our iPSC-derived samples as well as in comparison to previously published, publicly available data from other tissues and cell types. For our iPSC-derived samples, we used the R package edgeR (Robinson et al., 2010) to filter out lowly-expressed genes (counts per million (CPM) > 1 in 12/17 samples), resulting in 10,715 orthologous genes, and produced an MDS plot of our samples (Figure 1B). The greatest influence on our samples is species along PC1 and PC2, followed by separation of immature NPCs cells from mature cell types (neurons and astrocytes) along PC2 (Figure 1B). Notably, human samples were more variable than chimpanzee samples. One human cell line (H20961) showed significant variation across all cell types (SI Figures 2-4), however, the H20961 NPC sample was consistently an outlier, grouping outside of NPCs of either species, and was removed from subsequent analyses. There are no overt technical differences influencing this out-grouping (e.g. individual sex or age, RNA or cDNA library quality, read number, alignment percentages, SI Table 1). This cell line has successfully been used in before in other differentiation studies with no overtly different characteristics (Blake et al., 2018; Burrows et al., 2016; Eres et al., 2019; Pavlovic et al., 2018; I. G. Romero et al., 2015; Ward & Gilad, 2019; Ward et al., 2018).

To compare our samples to previously published data from cells and tissues, we downloaded raw RNA-Seq reads from the NCBI’s Gene Expression Omnibus (GEO) (Edgar, Domrachev, & Lash, 2002) and processed them from raw read counts through HT-Seq and orthologous gene matching in the same manner as our iPSC-derived samples. To compare our samples to those from primary neural cell types, we used RNA-Seq data from primary neurons and astrocytes obtained from four hippocampal astrocytes, four cortex astrocytes, and one cortical neuron from (Zhang et al., 2016) (GEO accession number GSE73721) and three pyramidal neuron samples (GEO accession numbers GSM2071331, GSM2071332, and GSM2071418) isolated from an unspecified brain region by the ENCODE project (Consortium, 2012; Davis et al., 2018). We also downloaded the tissue-level data from Brawand et al., (2011) (Brawand et al., 2011) from human and chimpanzee brain regions and non-neuronal tissue (heart, kidney, liver) (GEO accession number GSE30352) (SI Table 2 for details). Only genes with counts greater than zero in all samples were included (n=7,660) and were further filtered to include only those with CPM > 1 in all 23 samples (n=6,124). An MDS plot of normalized counts was generated using edgeR of the top 500 most differentially expressed genes in all samples (SI Figure 5).

### DIFFERENTIAL GENE EXPRESSION ANALYSES

In order to determine what genes were significantly differentially expressed in a species by cell type manner using, we used the R package edgeR’s (Robinson et al., 2010) generalize linear model (GLM) functionality with a design matrix accounting for an interaction between species (SP) and cell type (CT) (referred to as SPxCT DE analysis). We performed an analysis of variance (ANOVA)-like test for differences across all samples. Furthermore, in order to determine what differences existed between species for each cell type, we performed interspecies pairwise DE comparisons in a similar manner between NPCs, neurons, and astrocytes (referred to as CT-DE analyses). We also used the GLM for these analyses, but did not include more than one cell type in these analyses in order to include genes that may be cell-type specific. For all analyses, we used edgeR’s quasi-likelihood F-test and considered gene expression significantly different at a false discovery rate (FDR) of less than 5%. Normalization of data in edgeR for DE analyses ensured that DE is not dependent on original number of cells. All Venn diagrams were created using the R package Vennerable.

### CATEGORICAL ENRICHMENT ANALYSES

Uninformed pathway enrichment analyses were conducted using genes identified as differentially expressed from each DE comparison using gProfiler (Raudvere et al., 2019) with their functional enrichment tool (g:GOSt). Categorical enrichment analyses for overrepresented (enriched) and underrepresented (conserved) processes were conducted on all genes identified as differentially expressed (FDR < .05%) between species for individual cell types. Enrichments with a q-value of < .05 were considered significant.

### GENE SET ENRICHMENT ANALYSES

In order to investigate which metabolic pathways were enriched in a species’ CT, we used Gene Set Enrichment Analyses (GSEA) (Subramanian et al., 2005). We tested for enrichment of 23 *a priori* gene sets from the Molecular Signatures Database (MSigDB) (Liberzon et al., 2011) using the raw counts of the same set of genes used for the CT-DE pairwise comparisons. Gene sets were considered significantly enriched according to suggested thresholds (FDR < 25% and nominal p-value < .05) (Subramanian et al., 2005). Leading edge analyses determined a set of core enriched genes that most significantly influenced the enrichment of the gene set per phenotype.

### SELECTION ANALYSES

In order to determine if genes exhibiting significant interspecies differential expression also had evidence of positive selection in their coding sequences, we used nonsynonymous (dN) and synonymous (dS) nucleotide changes per gene for all genes expressed in iPSC-derived neural cells. These were obtained from Ensembl using biomaRt (Kinsella *et al*., 2011). These pre-calculated dN and dS values were originally computed by Ensembl using codeml and yn00 of the PAML package to compute dN and dS scores for each species in comparison to human (Herrero *et al*., 2016). A rate of change was calculated for each gene (dN/dS), where a dN/dS > 1 is indicative of positive selection (Herrero et al., 2016). In order to determine if there was evidence for noncoding selection, we analyzed promoter regions of genes involved in aerobic glycolysis. These genes were selected by downloading genelists from the Molecular Signatures Database (MSigDB) (Liberzon et al., 2011) for GO pathways involved in aerobic glycolysis (glycolysis, pyruvate conversion to lactate or acetyl CoA, TCA cycle, electron transport chain, oxidative phosphorylation) and further subset to those that were expressed in at least one sample (n=156). Signs of positive selection in non-coding regions adjacent to these genes were determined following the procedures outlined in Pizzollo et al., 2018 (Haygood et al., 2007; Pizzollo et al., 2018). Because these analyses are suited for nuclear encoded genes, we excluded mitochondrial-encoded genes (n=10). Rhesus macaque *(Macaca mulatta*) was used as an outgroup. After removing regions without sequences for all three species (human, chimpanzee, and rhesus macaque), we tested for positive selection in the human lineage of a total of 126 aerobic glycolysis genes.

### NETWORK SCHEMATIC

We constructed a focal set of signaling pathways based upon HumanCyc (P. Romero et al., 2005) in order to contextualize our DE results in the framework of a network signaling, and this is the diagram of the major pathways involved in aerobic glycolysis (glycolysis, pentose phosphate pathway (PPP), lactate conversion from pyruvate, and TCA cycle) shown in Figure 5. For each enzyme in the pathway, three blocks indicate expression of this enzyme in each cell type (left to right): NPCs, neurons, astrocytes. Color indicates level of expression (higher in human (red), higher in chimpanzee (blue), not expressed in this cell type (grey)).

## ACKNOWLEDGEMENTS

We would like to thank all members of the Babbitt laboratory for all of their support and feedback. We also thank Elena Vazey, Jason Kamilar, and Patricia Wadsworth for their insights and feedback.

## SUPPLEMENTAL FIGURES & TABLES

**SI Figure 1.**
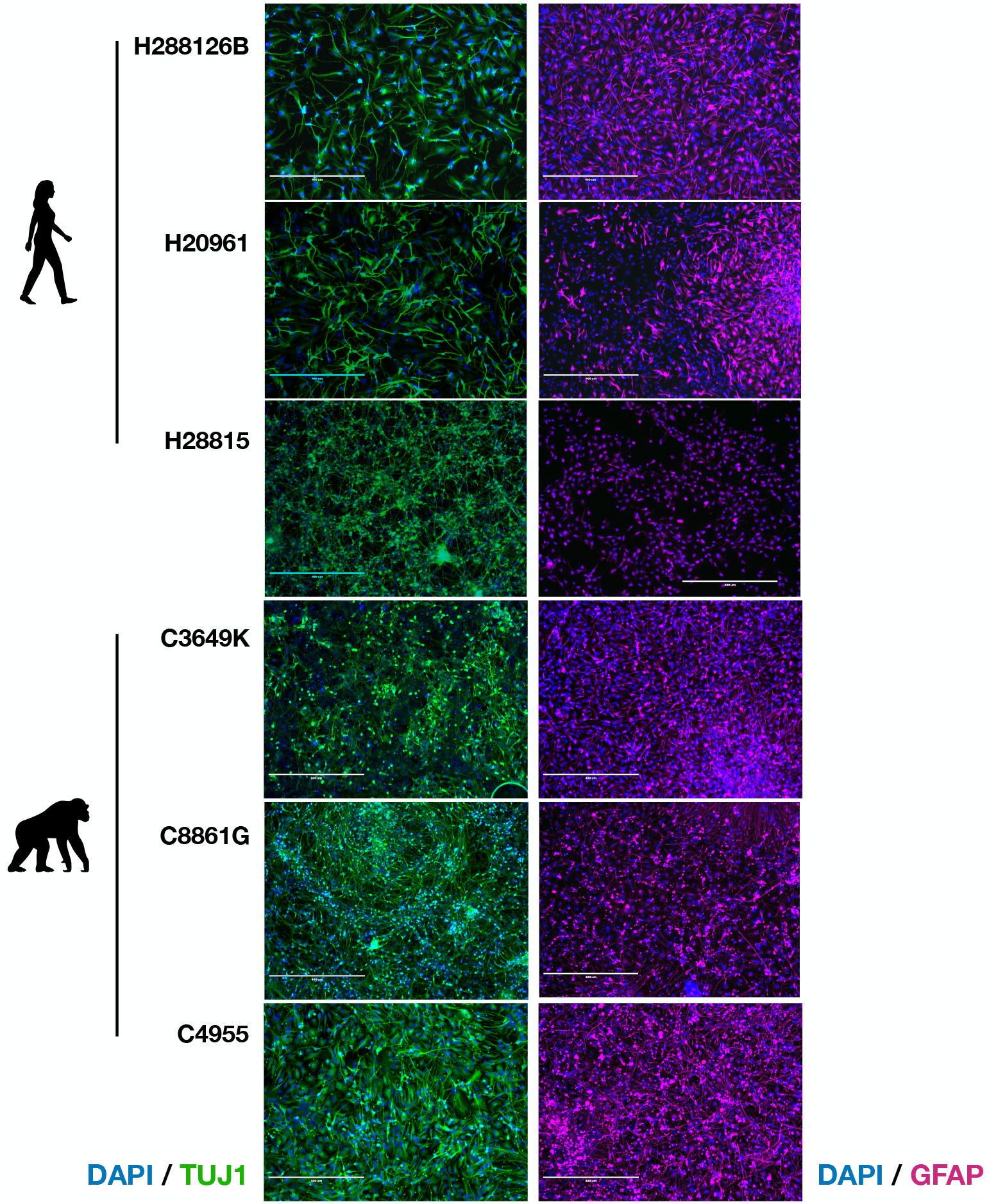
Differentiation and maturation of a human and chimpanzee iPSC lines into neural cell types. Immunofluorescent validation of matched-by-cell-line iPSC-derived neurons and astrocytes. Left column is images of cells immunofluorescently labeled for neuron-specific class III β-tubulin (TUJ1; Neuromics), and the right column is images of iPSC-derived astrocytes immunofluorescently labeled for GFAP (Sigma Aldrich), according to manufacturer’s suggestions. All cells for each cell type were harvested at similar timepoints: neurons at passage 3-4 and mature astrocytes at passage 5-6 post-differentiation from NPCs (SI Table 1).

**SI Figure 2.**
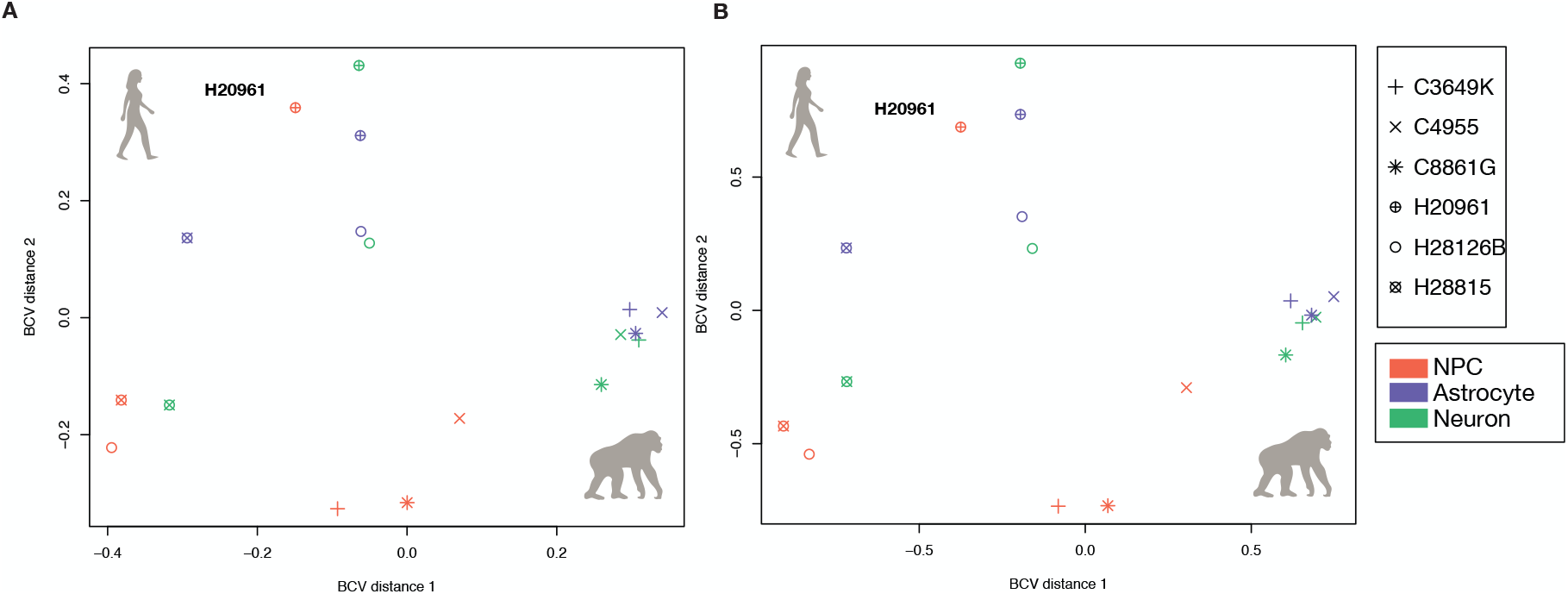
MDS plots of all iPSC-derived samples with shape indicating cell line. The same MDS plots for A) all genes expressed and B) the top 500 more differentially expressed genes as in main Figure 1C where shape indicates individual cell lines.

**SI Figure 3.**
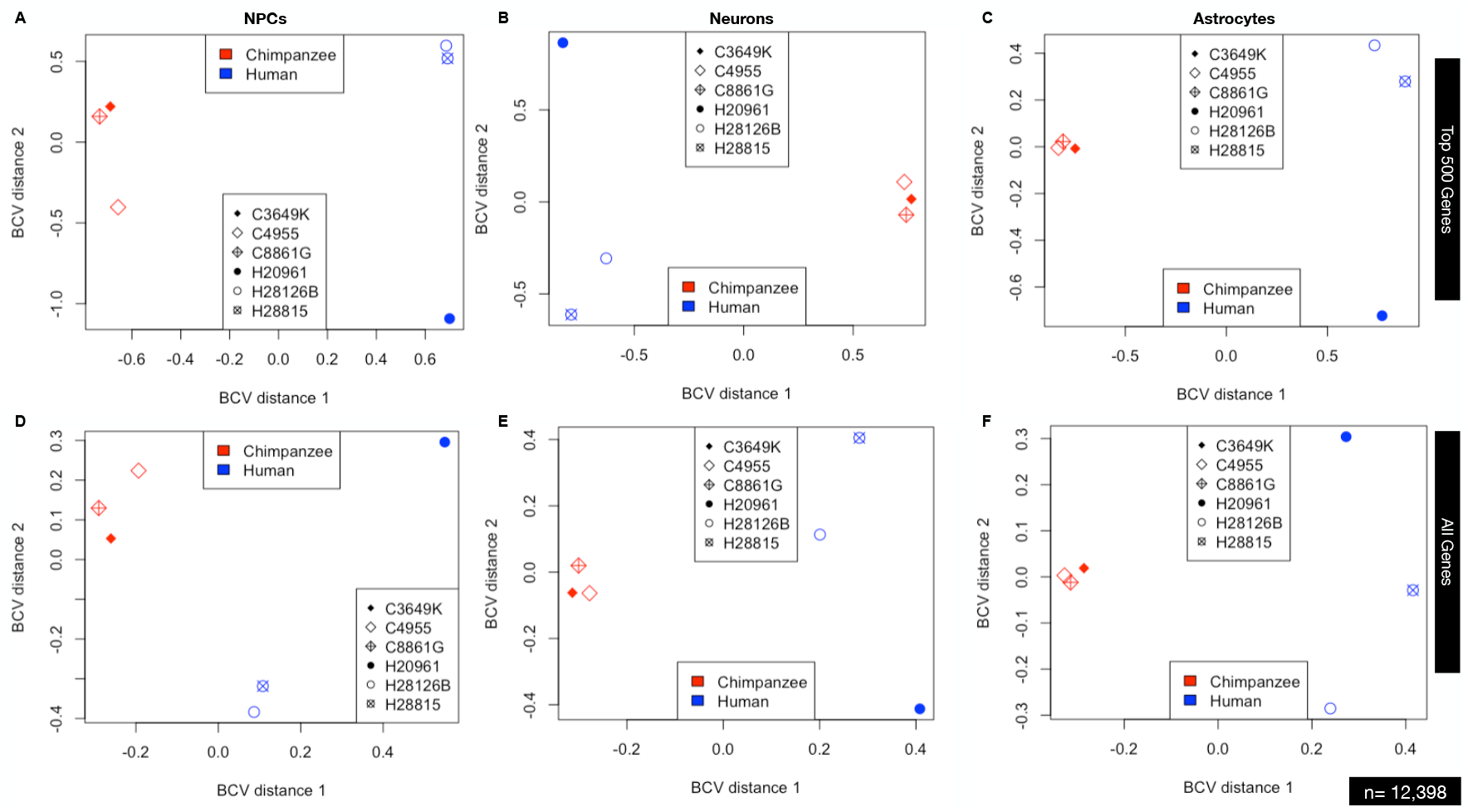
MDS plots of individual cell types (A & D – NPCs, B & E – neurons, C & F – astrocytes). Plots A-C are for the top 500 most differentially expressed genes while plots D-F are for all genes expressed.

**SI Figure 4.**
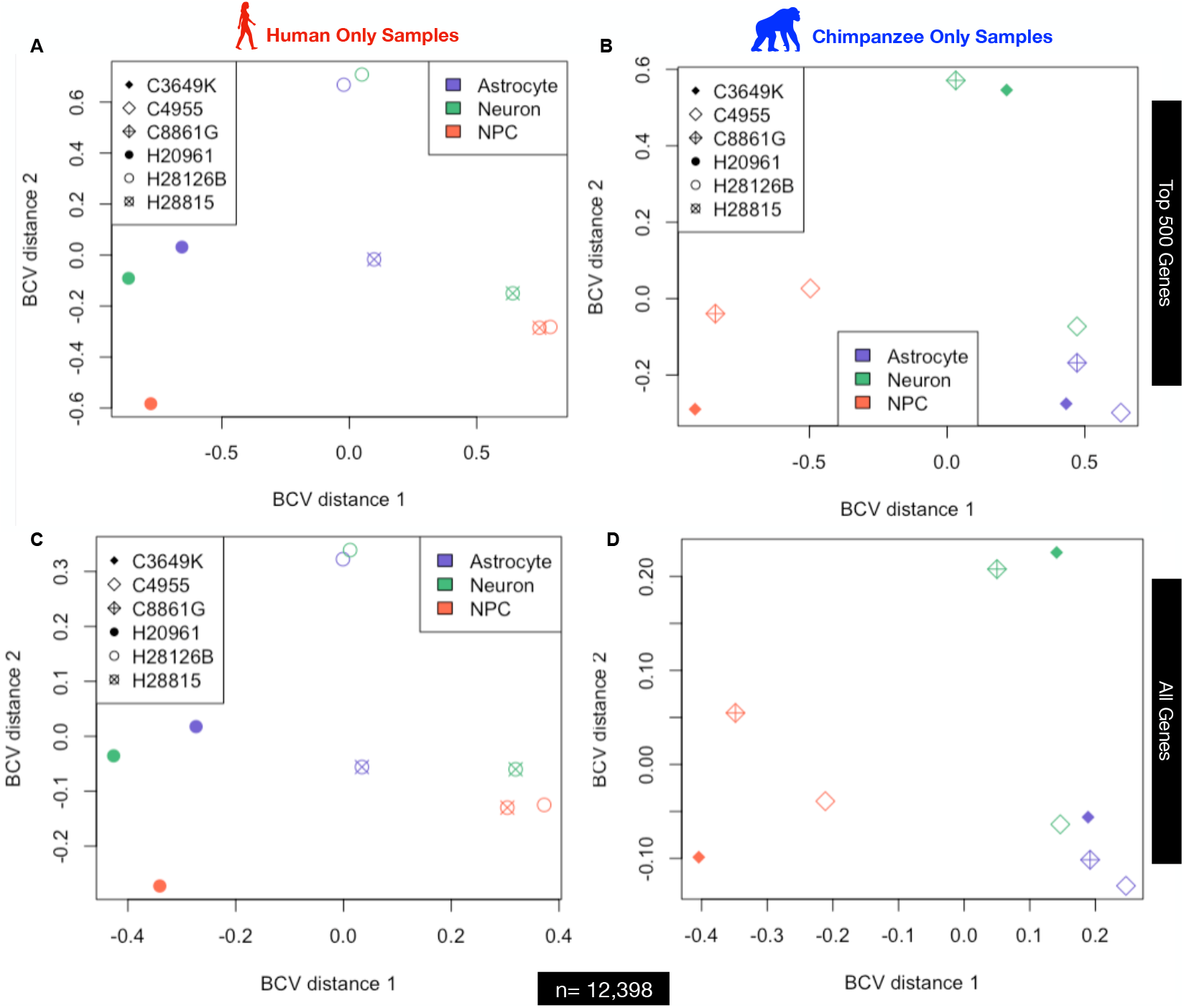
MDS plots of all A & C) human and B & D) chimpanzee samples by cell type. Plots A & B are for the top 500 most differentially expressed genes while plots C & D are for all genes expressed.

**SI Figure 5.**
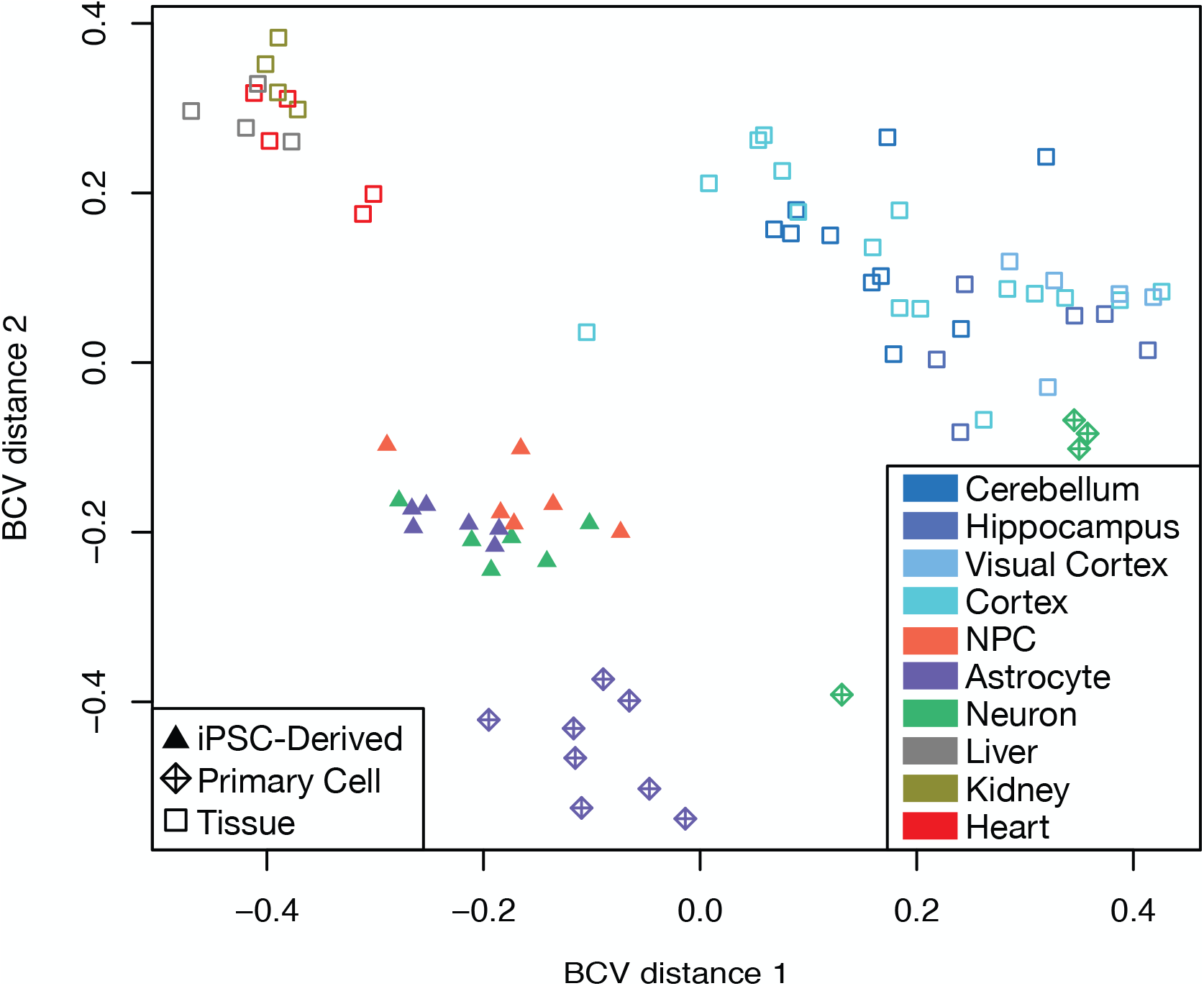
Human and chimpanzee iPSC-derived neural cells resemble primary neural cell types and tissue regions more than non-neuronal tissues. A PCoA of the iPSC-derived neural cells in comparison to whole-tissue RNA-Seq from four brain regions (cerebellum, hippocampus, prefrontal cortex, and visual cortex) from human and chimpanzee (3 individuals per species) (Babbitt et al, *in prep*.), brain and non-neuronal tissue from human and chimpanzee from (Brawand et al., 2011), as well as that from primary neurons and astrocytes (Consortium, 2012; Davis et al., 2018; Zhang et al., 2016) obtained from the Gene Expression Omnibus (GEO) (Edgar et al., 2002) Short Read Archive (SRA) (GEO accession numbers GSE30352, GSE73721, GSM2071331, GSM2071332, and GSM2071418). Label shape indicates the sample source tissue or cell type. Color refers to cell type or brain region.

**SI Figure 6.**
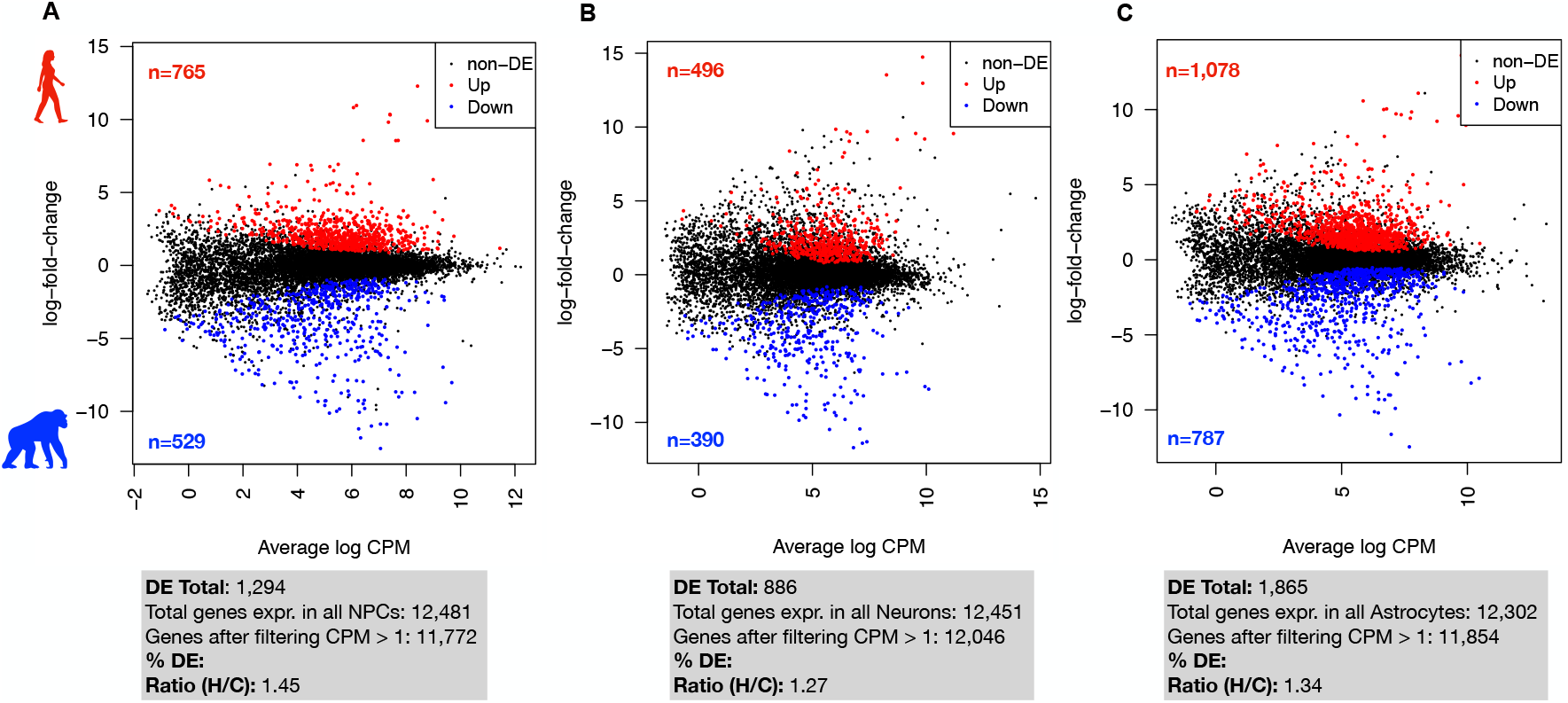
Distribution of differentially expressed genes between species for each cell type. MA plots of expression of all genes (average log CPM, x-axis) by their relative log fold-change (y-axis) from TopTags tables of pairwise, interspecies CT-DE comparisons made in edgeR for A) NPCs, B) neurons, and C) astrocytes. Color indicates differential expression status: black – non-DE, red – DE with higher expression in human, blue – DE with higher expression in chimpanzee). Number of genes identified as differentially expressed with higher expression in human and indicated in red text; for chimpanzee, in blue text.

**SI Figure 7.**
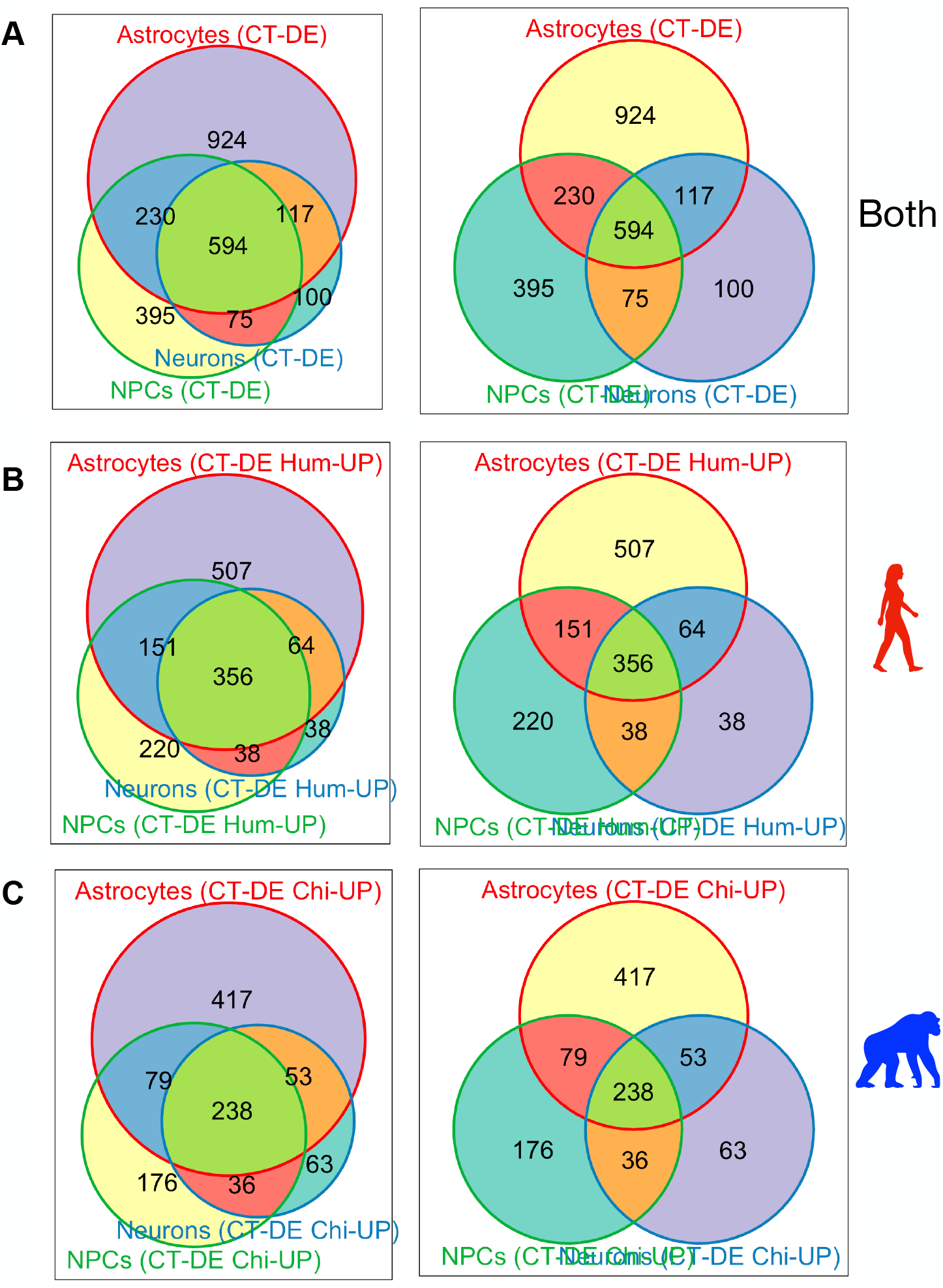
Overlap in interspecies CT-DE genes. Weighted (left column) and unweighted (right column) Venn diagrams made using R package Vennerable of overlap in genes per cell type exhibiting differential expression between species. Venn diagrams are for all CT-DE genes across all CT’s (A) and DE genes across all CT-DE comparisons with higher expression in human (B) or chimpanzee (C).

**SI Figure 8.**
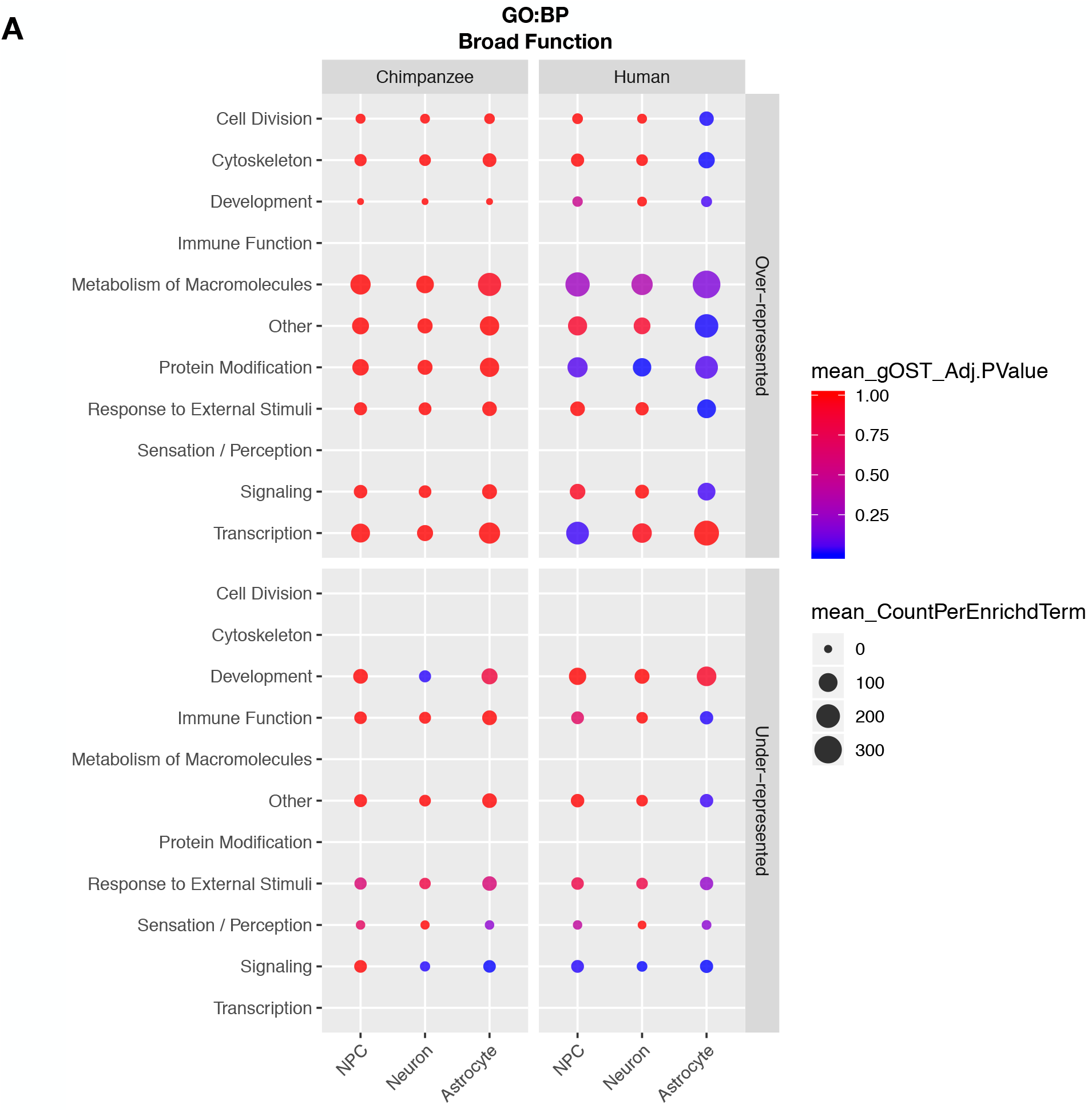
GO Biological Process (BP) enrichments. Plots of significantly over-represented (top panels) and under-represented (bottom panels) categories of GO BP terms determined by categorical enrichment analyses in genes with higher expression in chimpanzee (left panel) and human (right panel) for each cell type (x-axis). The categories (y-axis) represent groupings of multiple GO BP terms grouped by their general function. Size indicates the mean count and color indicates the mean adjusted enrichment p-value for all terms in that category.

**SI Figure 9.**
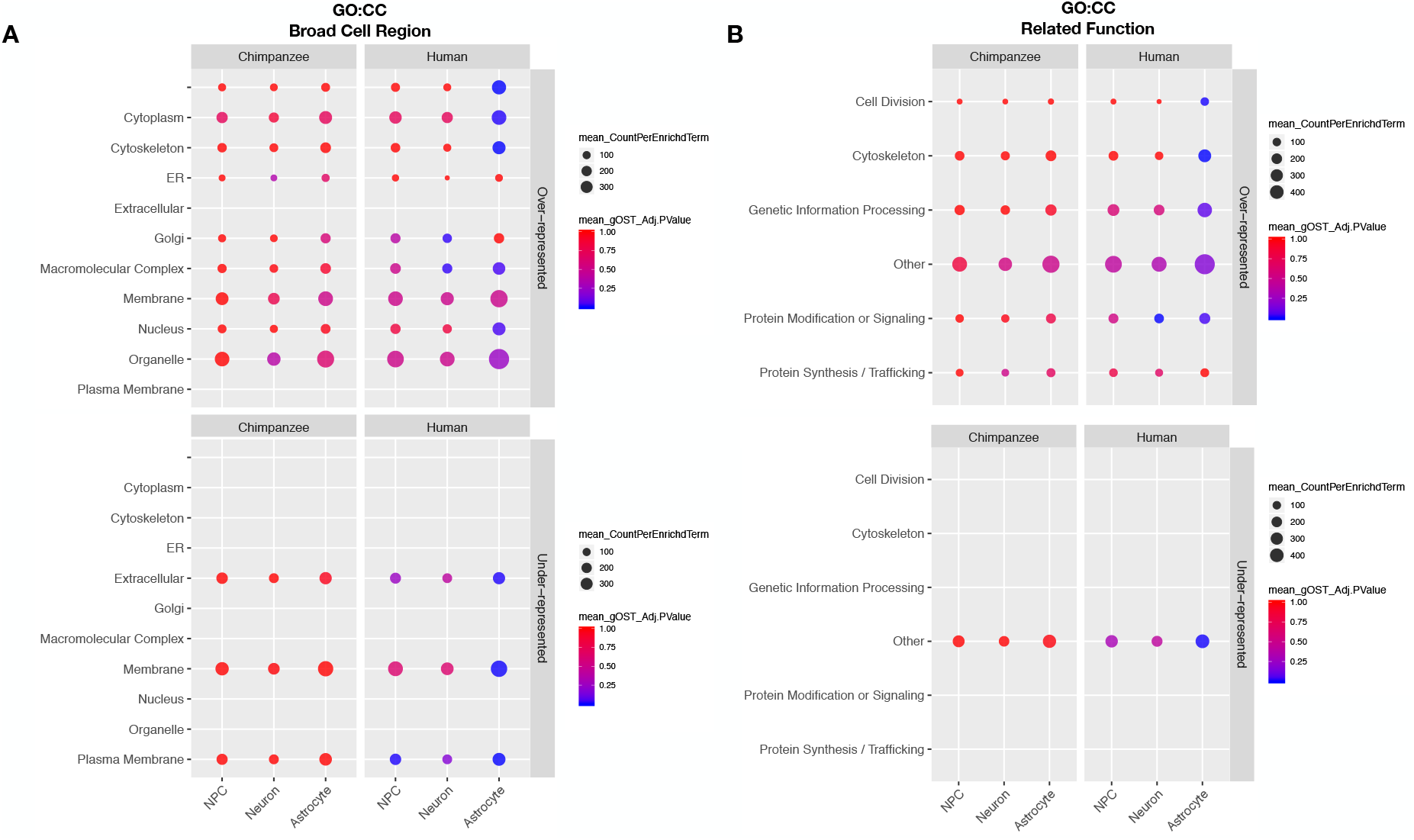
GO Cellular Component (CC) enrichments. Plots of significantly over-represented (top panels) and under-represented (bottom panels) categories of GO CC terms determined by categorical enrichment analyses in genes with higher expression in chimpanzee (left panel) and human (right panel) for each cell type (x-axis). The categories (y-axis) represent groupings of multiple GO CC terms grouped by their A) by their broad cell region and B) functions related to the cellular components enriched. Size indicates the mean count and color indicates the mean adjusted enrichment p-value for all terms in that category.

**SI Figure 10.**
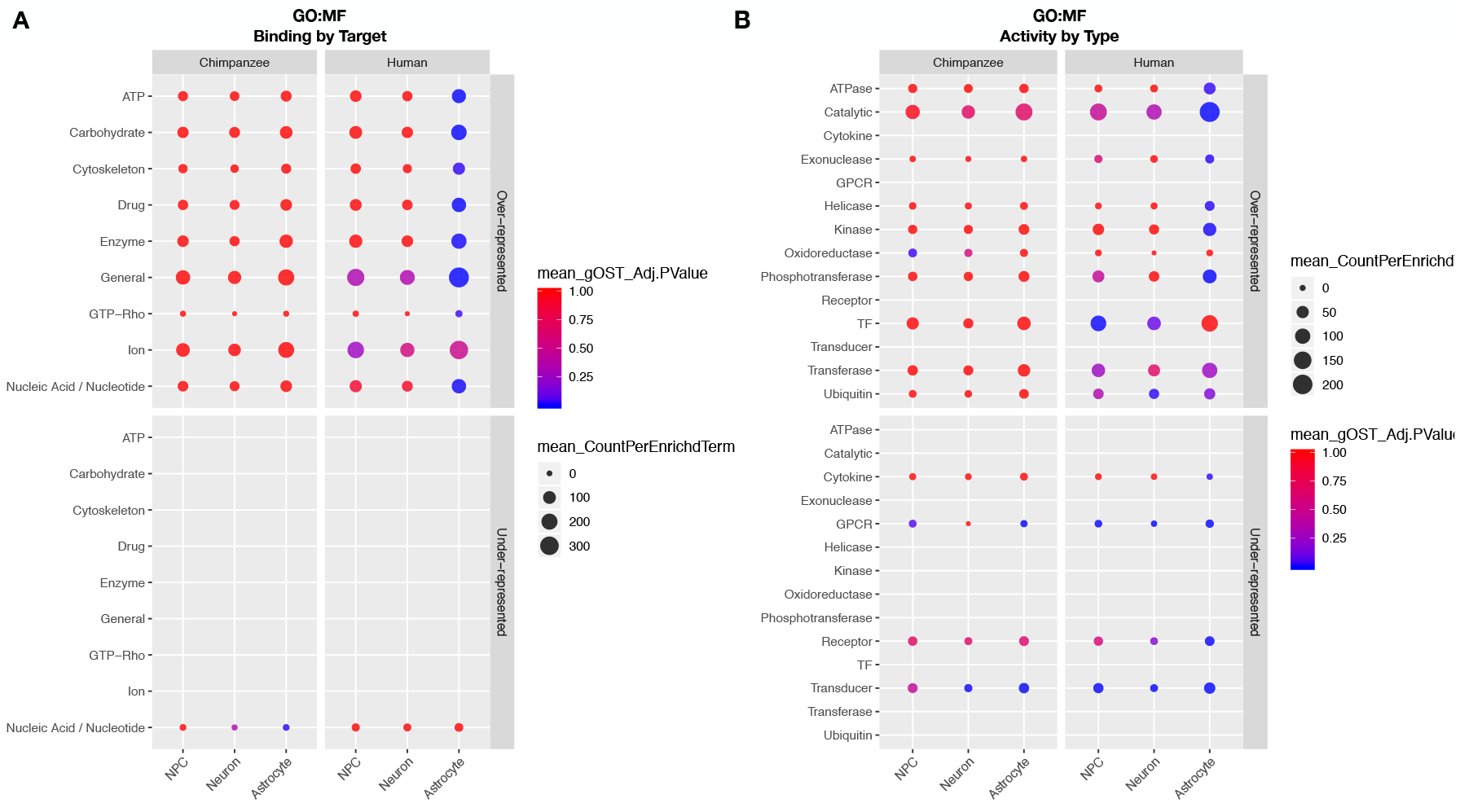
GO Molecular Function (MF) enrichments. Plots of significantly over-represented (top panels) and under-represented (bottom panels) categories of GO MF terms determined by categorical enrichment analyses in genes with higher expression in chimpanzee (left panel) and human (right panel) for each cell type (x-axis). The categories (y-axis) represent groupings of multiple GO MF terms grouped by their A) binding activity for particular substrates and B) specificity types of molecular activity. Size indicates the mean count and color indicates the mean adjusted enrichment p-value for all terms in that category.

**SI Figure 11.**
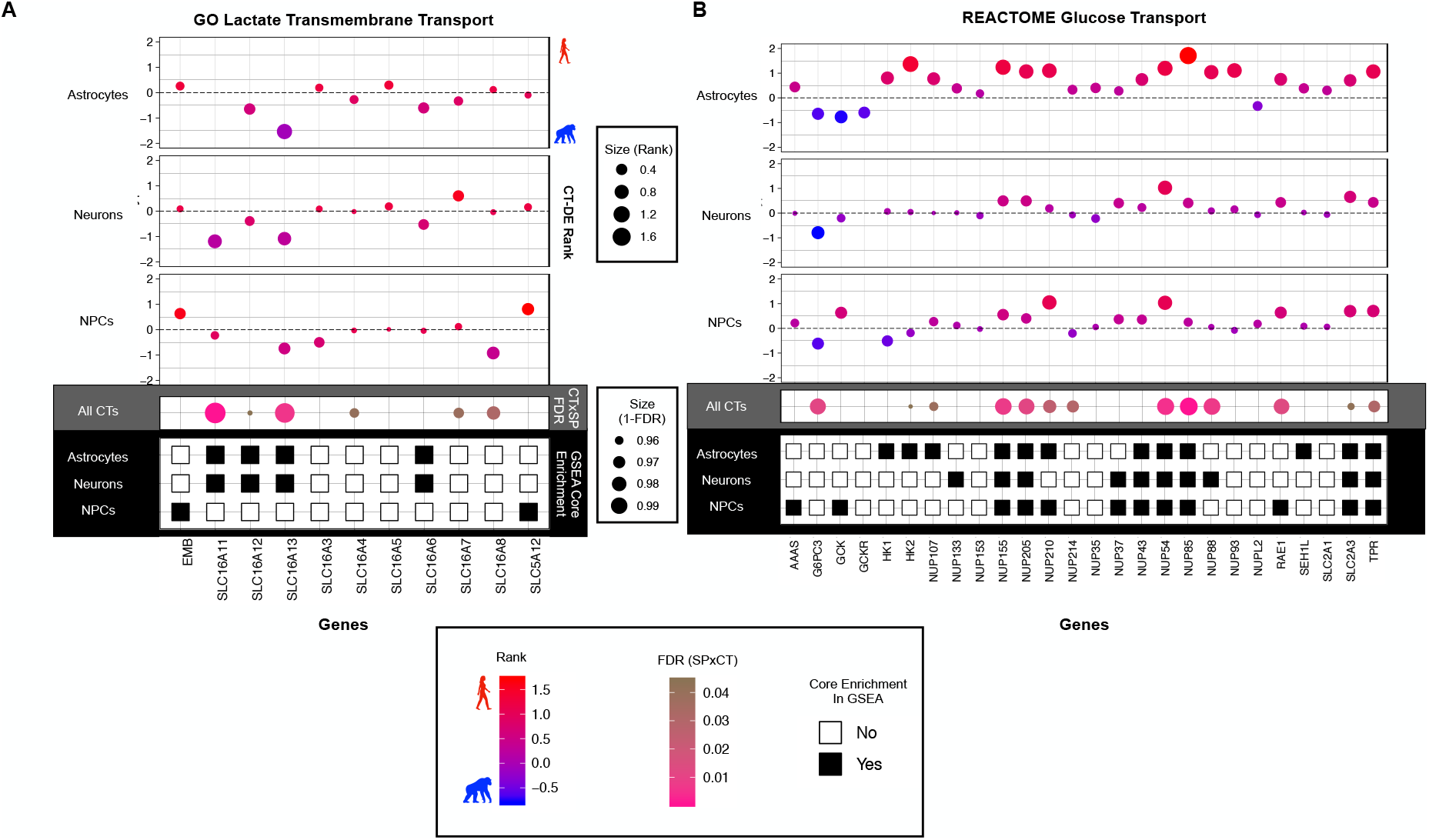
Neurons and astrocytes exhibit contrasting interspecies differences in lactate and glucose transport. Comparison of genes involved in the A) GO lactate transmembrane transport and B) REACTOME glucose transport gene sets determined as significantly enriched in one species by GSEA analyses. Each panel (white, grey, and black) indicates significance per gene for one of the functional enrichment analyses used (top/white - interspecies DE by cell type (CT-DE); middle/grey – SPxCT ANOVA-like DE, bottom/black – membership in core enrichment genes of leading edge GSEA analysis). (Top panel) Plot of CT-DE rank [(sign of logFC) x log10(FDR Q-value)] per each gene (x-axis),with values greater than zero indicating higher expression in human and values less than zero indicate higher expression in chimpanzee. Color spectrum and size also indicate rank (red – higher in human, blue – higher in chimpanzee, larger = higher rank). (Dark grey panel) Plot of FDR for each gene in the ANOVA-like SPxCT DE comparison, where color indicates significance (pink – lower/significant, brown – higher/non-significant) and size also indicates significance (1-FDR, larger = more significant). (Black panel) Plot of whether each gene was part of the core set of genes in GSEA leading edge analysis (black – yes, white – no). See SI Table 3 for DE results per gene and SI Table 6 for GSEA results per gene.

**SI Figure 12.**
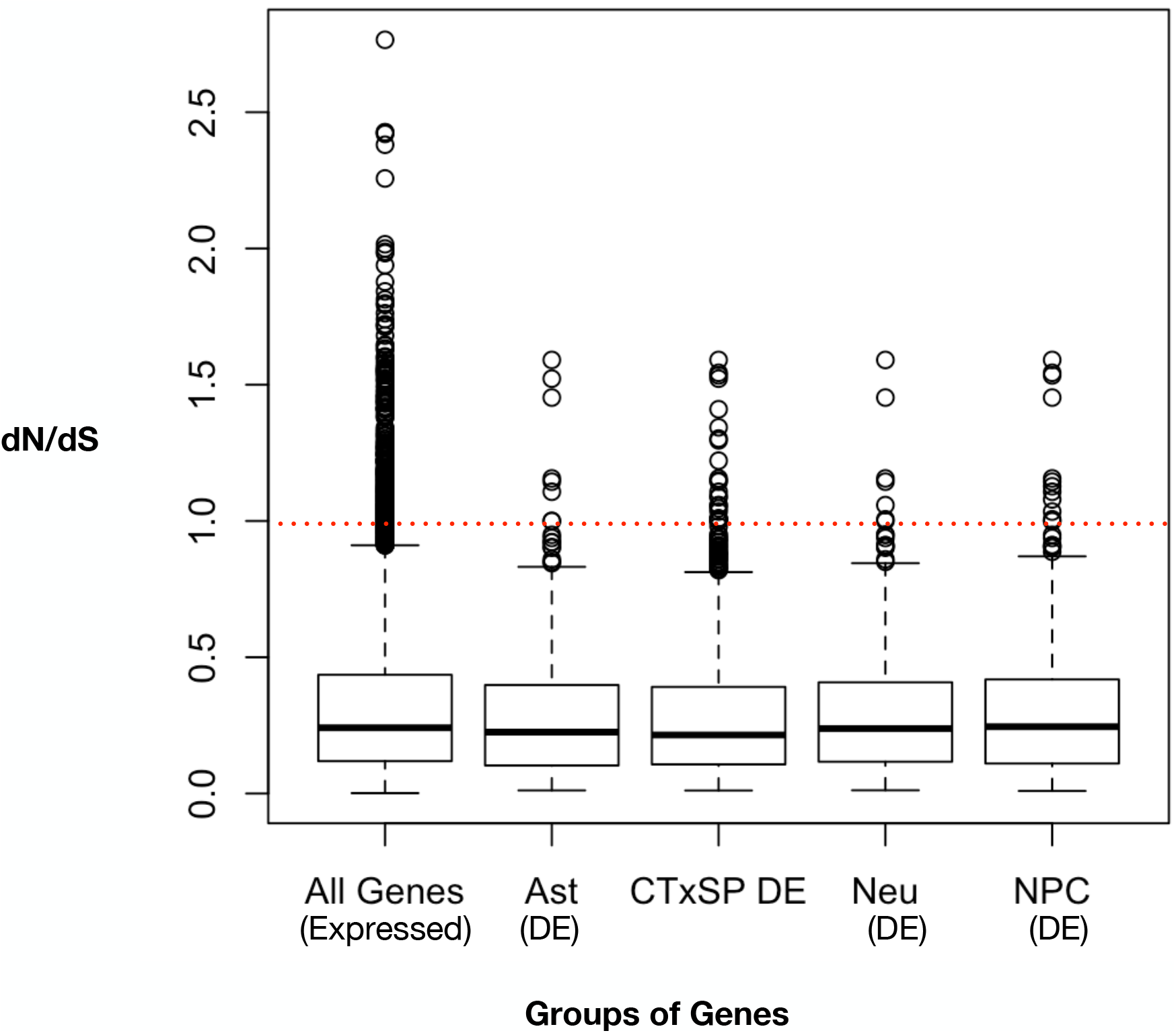
Positive selection in the coding regions of genes expressed in iPSC-derived neural cells. In order to determine if genes exhibiting significant interspecies differential expression also had evidence of positive selection in their coding sequences, we used nonsynonymous (dN) and synonymous (dS) nucleotide changes per gene for all genes expressed in iPSC-derived neural cells. A rate of change was calculated for each gene (dN/dS), where a dN/dS > 1 is indicative of positive selection.

**SI Figure 13.**
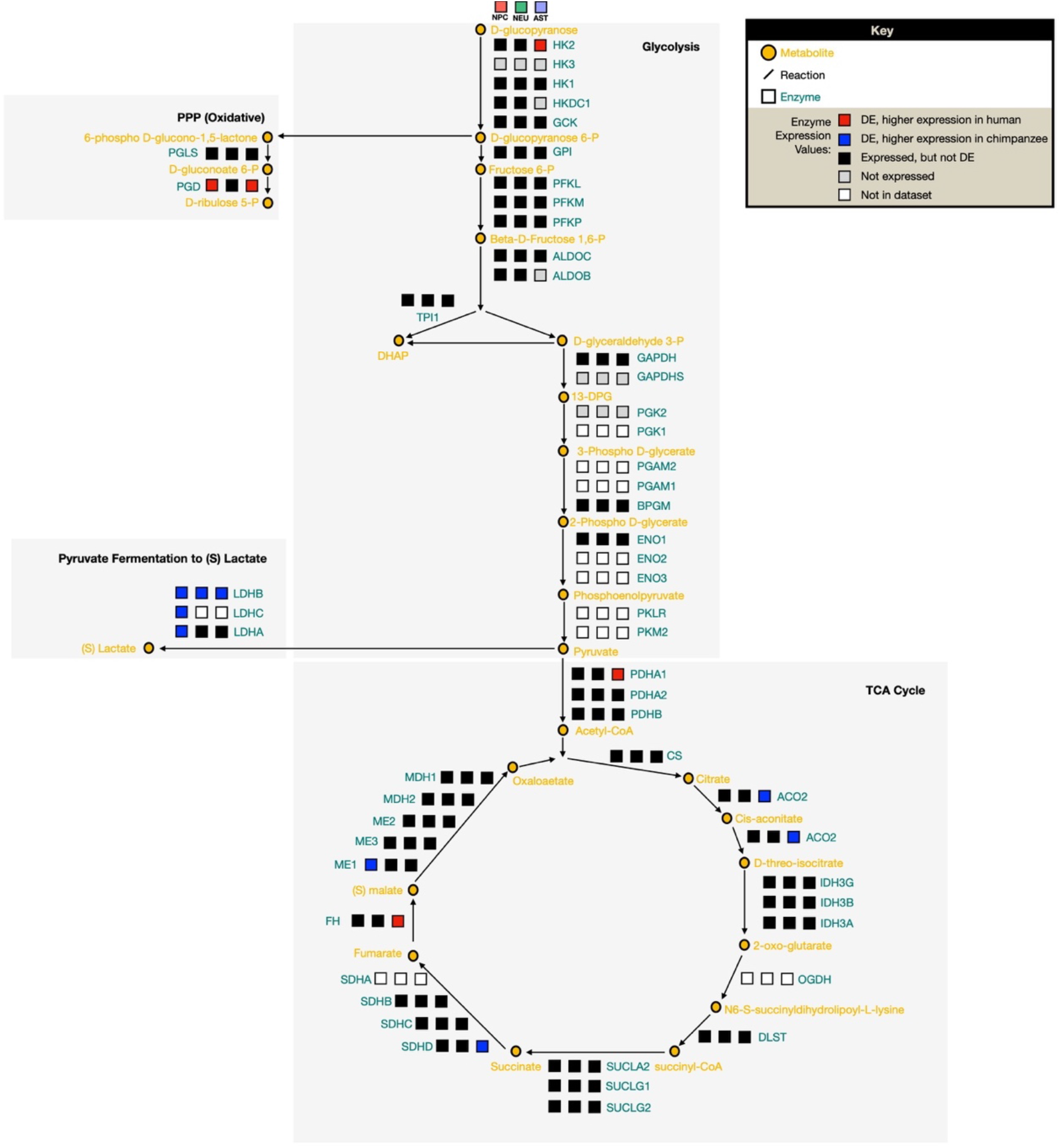
Full expression network of sub-pathways in aerobic glycolysis. We constructed a focal set of aerobic glycolysis signaling pathways in order to contextualize our DE results in the framework of a network signaling. A diagram of the major pathways involved in aerobic glycolysis (glycolysis, pentose phosphate pathway (PPP), lactate conversion from pyruvate, and TCA cycle). For each enzyme in the pathway, three blocks indicate expression of this enzyme in each cell type – left to right: NPCs, neurons, astrocytes. Color indicates level of expression (DE and higher in human (red), DE and higher in chimpanzee (blue), not expressed in this cell type (grey), expressed but not DE (black)).

## REFERENCES

Aiello, L. C., & Wheeler, P. (1995). The expensive-tissue hypothesis: the brain and the digestive system in human and primate evolution. Current anthropology, 36(2), 199–221.

Aken, B. L., Achuthan, P., Akanni, W., Amode, M. R., Bernsdorff, F., Bhai, J., … Clapham, P. (2016). Ensembl 2017. Nucleic Acids Research, 45(D1), D635–D642.

Almad, A. A., Doreswamy, A., Gross, S. K., Richard, J. P., Huo, Y., Haughey, N., & Maragakis, N. J. (2016). Connexin 43 in astrocytes contributes to motor neuron toxicity in amyotrophic lateral sclerosis. Glia, 64(7), 1154–1169.

Almeida, A., Moncada, S., & Bolaños, J. P. (2004). Nitric oxide switches on glycolysis through the AMP protein kinase and 6-phosphofructo-2-kinase pathway. Nature Cell Biology, 6(1), 45.

Amiri, A., Coppola, G., Scuderi, S., Wu, F., Roychowdhury, T., Liu, F., … Song, L. (2018). Transcriptome and epigenome landscape of human cortical development modeled in organoids. Science, 362(6420).

Anders, S., Pyl, P. T., & Huber, W. (2015). HTSeq—a Python framework to work with high-throughput sequencing data. Bioinformatics, 31(2), 166–169.

Antonazzo, G., Attrill, H., Brown, N., Marygold, S. J., McQuilton, P., Ponting, L., … Tweedie, S. (2017). Expansion of the Gene Ontology knowledgebase and resources. Nucleic Acids Research.

Ashburner, M., Ball, C. A., Blake, J. A., Botstein, D., Butler, H., Cherry, J. M., … Eppig, J. T. (2000). Gene Ontology: tool for the unification of biology. Nature Genetics, 25(1), 25.

Babbitt, C. C., Fedrigo, O., Pfefferle, A. D., Boyle, A. P., Horvath, J. E., Furey, T. S., & Wray, G. A. (2010). Both noncoding and protein-coding RNAs contribute to gene expression evolution in the primate brain. Genome Biology and Evolution, 2, 67–79.

Babbitt, C. C., Warner, L. R., Fedrigo, O., Wall, C. E., & Wray, G. A. (2011). Genomic signatures of diet-related shifts during human origins. Proceedings of the Royal Society B: Biological Sciences, 278(1708), 961–969.

Bakken, T. E., Miller, J. A., Luo, R., Bernard, A., Bennett, J. L., Lee, C.-K., … Sunkin, S. M. (2015). Spatiotemporal dynamics of the postnatal developing primate brain transcriptome. Human molecular genetics, 24(15), 4327–4339.

Bauernfeind, A. L., & Babbitt, C. C. (2014). The appropriation of glucose through primate neurodevelopment. Journal of Human Evolution, 77, 132–140.

Bauernfeind, A. L., Soderblom, E. J., Turner, M. E., Moseley, M. A., Ely, J. J., Hof, P. R., … Babbitt, C. C. (2015). Evolutionary divergence of gene and protein expression in the brains of humans and chimpanzees. Genome Biology and Evolution, 7(8), 2276–2288.

Blake, L. E., Thomas, S. M., Blischak, J. D., Hsiao, C. J., Chavarria, C., Myrthil, M., … Pavlovic, B. J. (2018). A comparative study of endoderm differentiation in humans and chimpanzees. Genome Biology, 19(1), 162.

Blekhman, R., Oshlack, A., Chabot, A. E., Smyth, G. K., & Gilad, Y. (2008). Gene regulation in primates evolves under tissue-specific selection pressures. PLoS Genetics, 4(11), e1000271.

Brawand, D., Soumillon, M., Necsulea, A., Julien, P., Csárdi, G., Harrigan, P., … Kircher, M. (2011). The evolution of gene expression levels in mammalian organs. Nature, 478(7369), 343.

Brown, A. M., Wender, R., & Ransom, B. R. (2001). Metabolic substrates other than glucose support axon function in central white matter. Journal of Neuroscience Research, 66(5), 839–843.

Brown, F., Harris, J., Leakey, R., & Walker, A. (1985). Early Homo erectus skeleton from west lake Turkana, Kenya. Nature, 316(6031), 788.

Burrows, C. K., Banovich, N. E., Pavlovic, B. J., Patterson, K., Romero, I. G., Pritchard, J. K., & Gilad, Y. (2016). Genetic variation, not cell type of origin, underlies the majority of identifiable regulatory differences in iPSCs. PLoS Genetics, 12(1), e1005793.

Camp, J. G., Badsha, F., Florio, M., Kanton, S., Gerber, T., Wilsch-Bräuninger, M., … Lancaster, M. (2015). Human cerebral organoids recapitulate gene expression programs of fetal neocortex development. Proceedings of the National Academy of Sciences, 112(51), 15672–15677.

Charnov, E. L., & Berrigan, D. (1993). Why do female primates have such long lifespans and so few babies? Or life in the slow lane. Evolutionary Anthropology, 1(6), 191–194.

Cho, I. K., Yang, B., Forest, C., Qian, L., & Chan, A. W. (2019). Amelioration of Huntington’s disease phenotype in astrocytes derived from iPSC-derived neural progenitor cells of Huntington’s disease monkeys. PLoS One, 14(3), e0214156.

Consortium, E. P. (2012). An integrated encyclopedia of DNA elements in the human genome. Nature, 489(7414), 57–74. doi:10.1038/nature11247

Davis, C. A., Hitz, B. C., Sloan, C. A., Chan, E. T., Davidson, J. M., Gabdank, I., … Cherry, J. M. (2018). The Encyclopedia of DNA elements (ENCODE): data portal update. Nucleic Acids Research, 46(D1), D794–D801. doi:10.1093/nar/gkx1081

di Domenico, A., Carola, G., Calatayud, C., Pons-Espinal, M., Muñoz, J. P., Richaud-Patin, Y., … Parameswaran, J. (2019). Patient-specific iPSC-derived astrocytes contribute to non-cell-autonomous neurodegeneration in Parkinson’s disease. Stem Cell Reports, 12(2), 213–229.

Diniz, L. P., Almeida, J. C., Tortelli, V., Lopes, C. V., Setti-Perdigão, P., Stipursky, J., … Alves-Leon, S. V. (2012). Astrocyte-induced synaptogenesis is mediated by transforming growth factor β signaling through modulation of D-serine levels in cerebral cortex neurons. Journal of Biological Chemistry, 287(49), 41432–41445.

Edgar, R., Domrachev, M., & Lash, A. E. (2002). Gene Expression Omnibus: NCBI gene expression and hybridization array data repository. Nucleic Acids Research, 30(1), 207–210. doi:10.1093/nar/30.1.207

Eres, I. E., Luo, K., Hsiao, C. J., Blake, L. E., & Gilad, Y. (2019). Reorganization of 3D genome structure may contribute to gene regulatory evolution in primates. PLoS Genetics, 15(7), e1008278.

Fabregat, A., Jupe, S., Matthews, L., Sidiropoulos, K., Gillespie, M., Garapati, P., … D’Eustachio, P. (2018). The Reactome Pathway Knowledgebase. Nucleic Acids Research, 46(D1), D649–D655. doi:10.1093/nar/gkx1132

Fagundes, N. J., Ray, N., Beaumont, M., Neuenschwander, S., Salzano, F. M., Bonatto, S. L., & Excoffier, L. (2007). Statistical evaluation of alternative models of human evolution. Proceedings of the National Academy of Sciences of the United States of America, 104(45), 17614–17619.

Goldberg, A., Wildman, D. E., Schmidt, T. R., Hüttemann, M., Goodman, M., Weiss, M. L., & Grossman, L. I. (2003). Adaptive evolution of cytochrome c oxidase subunit VIII in anthropoid primates. Proceedings of the National Academy of Sciences of the United States of America, 100(10), 5873–5878.

Grossman, L. I., Schmidt, T. R., Wildman, D. E., & Goodman, M. (2001). Molecular evolution of aerobic energy metabolism in primates. Molecular phylogenetics and evolution, 18(1), 26–36.

Grossman, L. I., Wildman, D. E., Schmidt, T. R., & Goodman, M. (2004). Accelerated evolution of the electron transport chain in anthropoid primates. TRENDS in Genetics, 20(11), 578–585.

Haygood, R., Babbitt, C. C., Fedrigo, O., & Wray, G. A. (2010). Contrasts between adaptive coding and noncoding changes during human evolution. Proceedings of the National Academy of Sciences of the United States of America, 200911249.

Haygood, R., Fedrigo, O., Hanson, B., Yokoyama, K.-D., & Wray, G. A. (2007). Promoter regions of many neural-and nutrition-related genes have experienced positive selection during human evolution. Nature Genetics, 39(9), 1140.

Herculano-Houzel, S. (2011). Scaling of brain metabolism with a fixed energy budget per neuron: implications for neuronal activity, plasticity and evolution. PLoS One, 6(3), e17514.

Herrero, J., Muffato, M., Beal, K., Fitzgerald, S., Gordon, L., Pignatelli, M., … Brent, S. (2016). Ensembl comparative genomics resources. Database, 2016.

Herrero-Mendez, A., Almeida, A., Fernández, E., Maestre, C., Moncada, S., & Bolaños, J. P. (2009). The bioenergetic and antioxidant status of neurons is controlled by continuous degradation of a key glycolytic enzyme by APC/C–Cdh1. Nature Cell Biology, 11(6), 747.

Hofman, M. A. (1983). Energy metabolism, brain size and longevity in mammals. The Quarterly Review of Biology, 58(4), 495–512.

Horvath, J. E., Ramachandran, G. L., Fedrigo, O., Nielsen, W. J., Babbitt, C. C., Clair, E. M. S., … Wall, C. E. (2014). Genetic comparisons yield insight into the evolution of enamel thickness during human evolution. Journal of Human Evolution, 73, 75–87.

Hüttemann, M., Helling, S., Sanderson, T. H., Sinkler, C., Samavati, L., Mahapatra, G., … Ramzan, R. (2012). Regulation of mitochondrial respiration and apoptosis through cell signaling: cytochrome c oxidase and cytochrome c in ischemia/reperfusion injury and inflammation. Biochimica et Biophysica Acta (BBA)-Bioenergetics, 1817(4), 598–609.

Kanton, S., Boyle, M. J., He, Z., Santel, M., Weigert, A., Sanchís-Calleja, F., … Han, D. (2019). Organoid single-cell genomic atlas uncovers human-specific features of brain development. Nature, 574(7778), 418–422.

Karbowski, J. (2007). Global and regional brain metabolic scaling and its functional consequences. BMC Biology, 5(1), 18.

Kersey, P. J., Allen, J. E., Allot, A., Barba, M., Boddu, S., Bolt, B. J., … Grabmueller, C. (2017). Ensembl Genomes 2018: an integrated omics infrastructure for non-vertebrate species. Nucleic Acids Research, 46(D1), D802–D808.

Khaitovich, P., Muetzel, B., She, X., Lachmann, M., Hellmann, I., Dietzsch, J., … Enard, W. (2004). Regional patterns of gene expression in human and chimpanzee brains. Genome Research, 14(8), 1462–1473.

Khrameeva, E., Kurochkin, I., Han, D., Guijarro, P., Kanton, S., Santel, M., … Sabirov, M. (2020). Single-cell-resolution transcriptome map of human, chimpanzee, bonobo, and macaque brains. Genome Research, 30(5), 776–789.

Kinsella, R. J., Kähäri, A., Haider, S., Zamora, J., Proctor, G., Spudich, G., … Kerhornou, A. (2011). Ensembl BioMarts: a hub for data retrieval across taxonomic space. Database, 2011.

Konopka, G., Friedrich, T., Davis-Turak, J., Winden, K., Oldham, M. C., Gao, F., … Preuss, T. M. (2012). Human-specific transcriptional networks in the brain. Neuron, 75(4), 601-617.

Kosiol, C., Vinař, T., da Fonseca, R. R., Hubisz, M. J., Bustamante, C. D., Nielsen, R., & Siepel, A. (2008). Patterns of positive selection in six mammalian genomes. PLoS Genetics, 4(8), e1000144.

Kuzawa, C. W., Chugani, H. T., Grossman, L. I., Lipovich, L., Muzik, O., Hof, P. R., … Lange, N. (2014). Metabolic costs and evolutionary implications of human brain development. Proceedings of the National Academy of Sciences, 111(36), 13010–13015.

Langmead, B., & Salzberg, S. L. (2012). Fast gapped-read alignment with Bowtie 2. Nature Methods, 9(4), 357.

Leonard, W. R., & Robertson, M. L. (1994). Evolutionary perspectives on human nutrition: the influence of brain and body size on diet and metabolism. American Journal of Human Biology, 6(1), 77–88.

Leonard, W. R., & Robertson, M. L. (1997). Comparative primate energetics and hominid evolution. American Journal of Physical Anthropology: The Official Publication of the American Association of Physical Anthropologists, 102(2), 265–281.

Leonard, W. R., Robertson, M. L., Snodgrass, J. J., & Kuzawa, C. W. (2003). Metabolic correlates of hominid brain evolution. Comparative Biochemistry and Physiology Part A: Molecular & Integrative Physiology, 136(1), 5–15.

Liberzon, A., Subramanian, A., Pinchback, R., Thorvaldsdóttir, H., Tamayo, P., & Mesirov, J. P. (2011). Molecular signatures database (MSigDB) 3.0. Bioinformatics, 27(12), 1739–1740.

Luo, C., Lancaster, M. A., Castanon, R., Nery, J. R., Knoblich, J. A., & Ecker, J. R. (2016). Cerebral organoids recapitulate epigenomic signatures of the human fetal brain. Cell Reports, 17(12), 3369–3384.

Mächler, P., Wyss, M. T., Elsayed, M., Stobart, J., Gutierrez, R., von Faber-Castell, A., … Romero-Gómez, I. (2016). In vivo evidence for a lactate gradient from astrocytes to neurons. Cell Metabolism, 23(1), 94–102.

Magistretti, P. J., & Allaman, I. (2015). A cellular perspective on brain energy metabolism and functional imaging. Neuron, 86(4), 883–901.

Martin, R. D. (1981). Relative brain size and basal metabolic rate in terrestrial vertebrates. Nature, 293(5827), 57.

Maxson, M. E., & Grinstein, S. (2014). The vacuolar-type H+-ATPase at a glance–more than a proton pump. In: The Company of Biologists Ltd.

McHenry, H. M. (1992). Body size and proportions in early hominids. American Journal of Physical Anthropology, 87(4), 407–431.

McHenry, H. M. (1994). Tempo and mode in human evolution. Proceedings of the National Academy of Sciences, 91(15), 6780–6786.

McKenzie, A. T., Wang, M., Hauberg, M. E., Fullard, J. F., Kozlenkov, A., Keenan, A., … Zhang, B. (2018). Brain Cell Type Specific Gene Expression and Co-expression Network Architectures. Scientific Reports, 8(1), 8868. doi:10.1038/s41598-018-27293-5

Meyer-Franke, A., Kaplan, M. R., Pfieger, F. W., & Barres, B. A. (1995). Characterization of the signaling interactions that promote the survival and growth of developing retinal ganglion cells in culture. Neuron, 15(4), 805–819.

Mink, J. W., Blumenschine, R. J., & Adams, D. B. (1981). Ratio of central nervous system to body metabolism in vertebrates: its constancy and functional basis. American Journal of Physiology-Regulatory, Integrative and Comparative Physiology, 241(3), R203–R212.

Muntané, G., Horvath, J. E., Hof, P. R., Ely, J. J., Hopkins, W. D., Raghanti, M. A., … Sherwood, C. C. (2014). Analysis of synaptic gene expression in the neocortex of primates reveals evolutionary changes in glutamatergic neurotransmission. Cerebral Cortex, 25(6), 1596–1607.

Nedergaard, M., Ransom, B., & Goldman, S. A. (2003). New roles for astrocytes: redefining the functional architecture of the brain. Trends in neurosciences, 26(10), 523–530.

Nelson, D. L., Lehninger, A. L., & Cox, M. M. (2008). Lehninger principles of biochemistry: Macmillan.

Ogata, H., Goto, S., Sato, K., Fujibuchi, W., Bono, H., & Kanehisa, M. (1999). KEGG: Kyoto encyclopedia of genes and genomes. Nucleic Acids Research, 27(1), 29–34.

Oldham, M. C., Horvath, S., & Geschwind, D. H. (2006). Conservation and evolution of gene coexpression networks in human and chimpanzee brains. Proceedings of the National Academy of Sciences of the United States of America, 103(47), 17973–17978.

Pamarthy, S., Kulshrestha, A., Katara, G. K., & Beaman, K. D. (2018). The curious case of vacuolar ATPase: regulation of signaling pathways. Molecular cancer, 17(1), 41.

Pavlovic, B. J., Blake, L. E., Roux, J., Chavarria, C., & Gilad, Y. (2018). A comparative assessment of human and chimpanzee iPSC-derived cardiomyocytes with primary heart tissues. Scientific Reports, 8(1), 15312.

Pellerin, L., & Magistretti, P. J. (1994). Glutamate uptake into astrocytes stimulates aerobic glycolysis: a mechanism coupling neuronal activity to glucose utilization. Proceedings of the National Academy of Sciences, 91(22), 10625–10629.

Penney, J., Ralvenius, W. T., & Tsai, L.-H. (2019). Modeling Alzheimer’s disease with iPSC-derived brain cells. Molecular psychiatry, 1–20.

Peters, C. R. (2007). Theoretical and actualistic ecobotanical perspectives on early hominin diets and paleoecology. Evolution of the human diet: the known, the unknown, and the unknowable, 233–261.

Pizzollo, J., Nielsen, W. J., Shibata, Y., Safi, A., Crawford, G. E., Wray, G. A., & Babbitt, C. C. (2018). Comparative serum challenges show divergent patterns of gene expression and open chromatin in human and chimpanzee. Genome Biology and Evolution, 10(3), 826–839.

Pond, S. L. K., Frost, S. D. W., & Muse, S. V. (2004). HyPhy: hypothesis testing using phylogenies. Bioinformatics, 21(5), 676–679. doi:10.1093/bioinformatics/bti079

Pontzer, H., Raichlen, D. A., Gordon, A. D., Schroepfer-Walker, K. K., Hare, B., O’Neill, M. C., … Isler, K. (2014). Primate energy expenditure and life history. Proceedings of the National Academy of Sciences of the United States of America, 111(4), 1433–1437.

Preuss, T. M. (2012). Human brain evolution: from gene discovery to phenotype discovery. Proceedings of the National Academy of Sciences, 109(Supplement 1), 10709–10716.

Raichle, M. E. (2010). Two views of brain function. Trends in cognitive sciences, 14(4), 180–190.

Raudvere, U., Kolberg, L., Kuzmin, I., Arak, T., Adler, P., Peterson, H., & Vilo, J. (2019). g:Profiler: a web server for functional enrichment analysis and conversions of gene lists (2019 update). Nucleic Acids Research, 47(W1), W191–W198. doi:10.1093/nar/gkz369

Reimand, J., Isserlin, R., Voisin, V., Kucera, M., Tannus-Lopes, C., Rostamianfar, A., … Bader, G. D. (2019). Pathway enrichment analysis and visualization of omics data using g:Profiler, GSEA, Cytoscape and EnrichmentMap. Nature Protocols, 14(2), 482–517. doi:10.1038/s41596-018-0103-9

Robinson, M. D., McCarthy, D. J., & Smyth, G. K. (2010). edgeR: a Bioconductor package for differential expression analysis of digital gene expression data. Bioinformatics, 26(1), 139–140.

Romero, I. G., Pavlovic, B. J., Hernando-Herraez, I., Zhou, X., Ward, M. C., Banovich, N. E., … Mitrano, A. (2015). A panel of induced pluripotent stem cells from chimpanzees: a resource for comparative functional genomics. Elife, 4, e07103.

Romero, P., Wagg, J., Green, M. L., Kaiser, D., Krummenacker, M., & Karp, P. D. (2005). Computational prediction of human metabolic pathways from the complete human genome. Genome Biology, 6(1), R2.

Schaffner, S. F., Foo, C., Gabriel, S., Reich, D., Daly, M. J., & Altshuler, D. (2005). Calibrating a coalescent simulation of human genome sequence variation. Genome Research, 15(11), 1576–1583.

Schneider, V. A., Graves-Lindsay, T., Howe, K., Bouk, N., Chen, H.-C., Kitts, P. A., … Albracht, D. (2017). Evaluation of GRCh38 and de novo haploid genome assemblies demonstrates the enduring quality of the reference assembly. Genome Research.

Shea, J. J. (2007). Lithic archaeology, or, what stone tools can (and can’t) tell us about early hominin diets. Evolution of the human diet: the known, the unknown and the unknowable, 321–351.

Snodgrass, J. J., Leonard, W. R., & Robertson, M. L. (2007). Primate bioenergetics: an evolutionary perspective. In Primate origins: adaptations and evolution (pp. 703–737): Springer.

Sonntag, K.-C., Ryu, W.-I., Amirault, K. M., Healy, R. A., Siegel, A. J., McPhie, D. L., … Cohen, B. M. (2017). Late-onset Alzheimer’s disease is associated with inherent changes in bioenergetics profiles. Scientific Reports, 7(1), 14038.

Stearns, S. (1992). The evolution of life histories. In: Oxford Univ. Press.

Stringer, C. B., & Andrews, P. (1988). Genetic and fossil evidence for the origin of modern humans. Science, 239(4845), 1263–1268.

Subramanian, A., Tamayo, P., Mootha, V. K., Mukherjee, S., Ebert, B. L., Gillette, M. A., … Lander, E. S. (2005). Gene set enrichment analysis: a knowledge-based approach for interpreting genome-wide expression profiles. Proceedings of the National Academy of Sciences, 102(43), 15545–15550.

Tekkök, S. B., Brown, A. M., Westenbroek, R., Pellerin, L., & Ransom, B. R. (2005). Transfer of glycogen-derived lactate from astrocytes to axons via specific monocarboxylate transporters supports mouse optic nerve activity. Journal of Neuroscience Research, 81(5), 644–652.

Uddin, M., Goodman, M., Erez, O., Romero, R., Liu, G., Islam, M., … Wildman, D. E. (2008). Distinct genomic signatures of adaptation in pre-and postnatal environments during human evolution. Proceedings of the National Academy of Sciences, 105(9), 3215–3220.

Vander Heiden, M. G., Cantley, L. C., & Thompson, C. B. (2009). Understanding the Warburg effect: the metabolic requirements of cell proliferation. Science, 324(5930), 1029–1033.

Vander Heiden, M. G., Locasale, J. W., Swanson, K. D., Sharfi, H., Heffron, G. J., Amador-Noguez, D., … Asara, J. M. (2010). Evidence for an alternative glycolytic pathway in rapidly proliferating cells. Science, 329(5998), 1492–1499.

Varki, A., & Altheide, T. K. (2005). Comparing the human and chimpanzee genomes: searching for needles in a haystack. Genome Research, 15(12), 1746–1758.

Varki, N. M., & Varki, A. (2015). On the apparent rarity of epithelial cancers in captive chimpanzees. Philosophical Transactions of the Royal Society B: Biological Sciences, 370(1673), 20140225.

Velasco, S., Kedaigle, A. J., Simmons, S. K., Nash, A., Rocha, M., Quadrato, G., … Regev, A. (2019). Individual brain organoids reproducibly form cell diversity of the human cerebral cortex. Nature, 570(7762), 523–527.

Volkenhoff, A., Weiler, A., Letzel, M., Stehling, M., Klämbt, C., & Schirmeier, S. (2015). Glial glycolysis is essential for neuronal survival in Drosophila. Cell Metabolism, 22(3), 437–447.

Ward, M. C., & Gilad, Y. (2019). A generally conserved response to hypoxia in iPSC-derived cardiomyocytes from humans and chimpanzees. eLife, 8, e42374.

Ward, M. C., Zhao, S., Luo, K., Pavlovic, B. J., Karimi, M. M., Stephens, M., & Gilad, Y. (2018). Silencing of transposable elements may not be a major driver of regulatory evolution in primate iPSCs. Elife, 7, e33084.

West, G. B., Brown, J. H., & Enquist, B. J. (2001). A general model for ontogenetic growth. Nature, 413(6856), 628.

Wildman, D. E., Wu, W., Goodman, M., & Grossman, L. I. (2002). Episodic positive selection in ape cytochrome c oxidase subunit IV. Molecular Biology and Evolution, 19(10), 1812–1815.

Wu, W., Goodman, M., Lomax, M. I., & Grossman, L. I. (1997). Molecular evolution of cytochrome c oxidase subunit IV: evidence for positive selection in simian primates. Journal of molecular evolution, 44(5), 477–491.

Yang, Z. (2007). PAML 4: phylogenetic analysis by maximum likelihood. Molecular Biology and Evolution, 24(8), 1586–1591.

Yu, Y., Karbowski, J., Sachdev, R. N., & Feng, J. (2014). Effect of temperature and glia in brain size enlargement and origin of allometric body-brain size scaling in vertebrates. BMC Evolutionary Biology, 14(1), 178.

Zhang, Y., Sloan, S. A., Clarke, L. E., Caneda, C., Plaza, C. A., Blumenthal, P. D., … Li, G. (2016). Purification and characterization of progenitor and mature human astrocytes reveals transcriptional and functional differences with mouse. Neuron, 89(1), 37–53.

Zhao, J., Davis, M. D., Martens, Y. A., Shinohara, M., Graff-Radford, N. R., Younkin, S. G., … Bu, G. (2017). APOE ε4/ε4 diminishes neurotrophic function of human iPSC-derived astrocytes. Human molecular genetics, 26(14), 2690–2700.

Zheng, X., Boyer, L., Jin, M., Mertens, J., Kim, Y., Ma, L., … Hunter, T. (2016). Metabolic reprogramming during neuronal differentiation from aerobic glycolysis to neuronal oxidative phosphorylation. Elife, 5, e13374.

